# Neuroanatomy of the Accessory Olfactory Bulb in the Fossorial Water Vole

**DOI:** 10.1101/2024.10.11.617790

**Authors:** Sara Ruiz-Rubio, Irene Ortiz-Leal, Mateo V. Torres, Mostafa G. A. Elsayed, Aitor Somoano, Pablo Sanchez-Quinteiro

## Abstract

The accessory olfactory bulb (AOB) plays a key role in processing chemical signals crucial for species-specific social and reproductive behaviors. While extensive research has focused on the vomeronasal system of laboratory rodents, less is known about wild species, particularly those that rely heavily on chemical communication. This study aims to characterize the morphological and neurochemical organization of the AOB in the fossorial water vole (*Arvicola scherman*), a subterranean rodent species from the family Cricetidae. We have employed histological techniques, including Nissl and hematoxylin staining, as well as immunohistochemical and lectin-histochemical markers, to assess the AOB structure. Our findings reveal that the AOB of the water vole exhibits a distinct laminar organization with prominent mitral cells in the mitral-plexiform layer, as well as dense labeling of periglomerular and short-axon cells in the glomerular layer. Lectin histochemistry further confirmed zonation patterns analogous to those seen in other rodent species. Immunohistochemical analysis demonstrated significant expression of PGP 9.5, suggesting its involvement in maintaining neuronal activity within the AOB. In contrast, the absence of SMI-32 labeling in the AOB, compared to its strong expression in the main olfactory bulb, highlights functional distinctions between these two olfactory subsystems. These structural and neurochemical characteristics suggest that the AOB of the fossorial water vole is adapted for enhanced processing of chemosensory signals, which may play a pivotal role in its subterranean lifestyle. Our results provide a foundation for future studies exploring the functional implications of these adaptations, including potential improvements in the integrated management of these vole populations.

## Introduction

In the past, olfaction was considered a singular system specialized in detecting volatile odor compounds (Bembibre and Strlič 2022). Today, it is known that the olfactory system comprises several subsystems, each with distinct anatomical bases and functions (Menini 2010; Barrios et al. 2014b), and there is also significant cooperation among these subsystems (Salazar et al. 2016). Among them, the vomeronasal system (VNS) stands out, responsible for chemical communication, namely the detection of a diverse range of chemical signals that mediate innate responses crucial for the coordination of species-specific social and reproductive behaviors (Halpern and Martínez-Marcos 2003; Yohe and Krell 2023). Among these chemical signals, pheromones are prominent as molecules that trigger innate intraspecific behaviors, primarily sexual and reproductive behaviors. Kairomones also play a critical role, producing an interspecific aversive effect (Ortiz-Leal et al. 2020). Additionally, the detection of major histocompatibility complex molecules by the VNS is crucial, as they guide individuals in mating decisions (Scott 2003; Overath et al. 2014).

The VNS comprises a sensory receptor structure called the vomeronasal organ (VNO), which is internally lined by a neuroepithelium analogous to that of the olfactory system, yet distinct in its characteristics (Salazar et al. 1997; Salazar et al. 2003; Kondoh et al. 2022). Vomeronasal receptors, initially identified as two major families, V1R and V2R (Dulac and Axel 1995; Ryba and Tirindelli 1997), are located on the luminal surface of the epithelium, although other types of neuroreceptors, such as formyl peptide receptors, have been characterized subsequently (Boillat et al. 2021). The neuroepithelium transmits information to the central nervous system via the vomeronasal nerves, specifically to a distinct area within the olfactory bulb, known as the accessory olfactory bulb (AOB) (Schröder et al. 2020; Torres et al. 2022). The presence of these neural projections is crucial for identifying the AOB, as this structure exhibits considerable morphological diversity among mammals (Meisami and Bhatnagar 1998). Finally, the secondary projections of the VNS, predominantly to the vomeronasal amygdala, were demonstrated thanks to comprehensive tracer studies conducted in the 1970s (Winans and Scalia 1970; Scalia and Winans 1975). The AOB is the main integrative neural center for the information detected by the VNO, as most of the sensory information collected by the VNO passes through the AOB (Holy 2018). However, it has been postulated that in rodents and lagomorphs, as well as in other groups of mammals, such as canids, there exists a projection of vomeronasal information to the transitional area between the main olfactory bulb (MOB) and the AOB, a region known as the olfactory limbus (Ortiz-Leal et al. 2023). Additionally, the AOB in mammals presents a broad diversity, both from a macroscopic and microscopic anatomical standpoint (Switzer, III. et al. 1980; Fernández-Aburto et al. 2020). This diversity even requires, in certain mammalian groups, the use of histochemical and neurochemical techniques to precisely determine the presence of an AOB. This is the case with canids (Nakajima et al. 1998; Torres et al. 2022), mustelids (Kelliher et al. 2001), or even humans (Chuah and Zheng 1987), for whom there are few available descriptions. In other species, however, the AOB occupies a very noticeable volume in relation to the main olfactory bulb. This is the case for rodents (Mucignat 2004), marsupials (Schneider et al. 2012), and lagomorphs (Villamayor et al. 2020), whose AOB is macroscopically evident. Ultimately, these differences imply that when characterizing the AOB of a species, extrapolations should be avoided, due to the striking differences, even among closely related mammalian families, such as those in the order Chiroptera (Frahm and Bhatnagar 1980). This fact becomes even more evident when considering microscopic organization. Although, generally speaking, the AOB has a laminar structure analogous to that of the MOB, both the lamination and the pattern of glomerular organization, as well as the cellularity of each layer, vary significantly depending on the species (Chengetanai et al. 2020; Ortiz-Leal et al. 2024). Likewise, striking differences have been described in the organization of the projection cells, the mitral cells, which exhibit considerable diversity in their morphological and topographic patterns depending on the animal group.

Given the importance of chemical communication in rodents and the fact that rats and mice have been the most utilized and studied models in the characterization of the mammalian nervous system, there is extensive information regarding the AOB in laboratory rodents (Larriva-Sahd 2008; Martín-López et al. 2012). However, despite the large number of families that include the order of rodents, very few of them have been studied in terms of the microscopic and neurochemical organization of the AOB. To our knowledge, the AOB of wild rodents has only been studied in the capybara *Hydrochoerus hydrochaeris* (Suárez et al. 2011b; Torres et al. 2020), the degu, *Octodon degus* (Suárez and Mpodozis 2009; Suárez et al. 2011a), the ground squirrel, *Spermophilus beecheyi* (Suárez et al. 2011a) and the beaver, *Castor fiber* (Tomiyasu et al. 2022), species belonging respectively to the Caviidae, Octodontidae, Sciuridae, and Castoridae families, and their AOBs show notorious differences compared to those of laboratory rodents. In this study, we have focused in the morphological and neurochemical characterization of the AOB of a rodent belonging to the Family Cricetidae, subfamily Arvicolinae, specifically the fossorial water vole, *Arvicola scherman* (formerly *Arvicola terrestris*, (Balmori-de La Puente et al. 2022)), of which information available on its olfactory subsystems is limited to the VNO (Ruiz-Rubio et al. 2024). As for the AOB of the Cricetids, to our knowledge information only exists about the hamster *Mesocricetus auratus*, a domestic species of the subfamily Cricetinae (Taniguchi et al. 1993b; Nakajima et al. 1996; Nakajima et al. 1998; Saito et al. 1999).

Given that the fossorial water vole lives mainly underground in extensive burrow systems (Airoldi 1976), it relies more on chemical senses than on physical senses such as hearing and vision (Dennis et al. 2020). Indeed, the comprehensive study of the species VNO, both morphologically and neurochemically (Ruiz-Rubio et al. 2024), represents a first step towards confirming the importance of kairomone and pheromone detection. The study of the organization of the chemosensory systems of *Arvicola scherman* is of particular interest because it is a species that can spread and reach population of up to 1000 voles/ha during population peaks in agroecosystems in the main mountainous areas of Europe, being considered a serious agricultural pest in much of its range (Somoano 2020). This species is also considered a human health hazard serving as a reservoir for zoonotic pathogens like *Leptospira* spp. (Giraudoux et al. 2009) or *Borrelia burgdorferi* s.l. (Espí et al. 2017), and parasites such as *Toxoplasma gondii* (Fuehrer et al. 2010) or *Echinococcus multilocularis* (Robardet et al. 2011).

Kairomones are naturally released by predators and detected by their prey (Sbarbati and Osculati 2006; Fortes-Marco et al. 2013), causing an aversive reaction based on physiological changes such as an increase in blood cortisol levels, systemic stress, and notable behavioral alterations, such as a change in habitat use (Dielenberg and McGregor 2001; Horii et al. 2010; Takahashi 2014). This paves the way for the use of synthetic kairomone and pheromone analogs that could be employed in chemical-based strategies to improve integrated pest management in fossorial water voles (Apfelbach et al. 2005; Papes et al. 2010). An example of the feasibility of this approach is illustrated by the field study conducted by Poissenot et al. (2023a; 2023b), based on the use of volatile organic compounds present in the urine and in the lateral scent gland (LSG) of conspecifics (Nagnan-Le Meillour et al. 2019). The success of such strategies underscores the necessity for in-depth neuroanatomical and neurochemical understanding of the species, enabling more effective targeting and application of these compounds

In this work we address the neurochemical study of the AOB of the water vole using macroscopic and microscopic dissection techniques, conventional histological methods, primarily Nissl and hematoxylin staining, as well as immunohistochemical and lectin-histochemical techniques. This will provide the basis for further research to assess the physiological impact of kairomones on individual organisms, which could ultimately improve integrated management of overabundant vole populations

## Methods

In total, 10 specimens of *Arvicola scherman*, with an equal representation of both sexes, were obtained from grasslands in the village of O Biduedo, Municipality of Triacastela. The animals were caught using snap traps (Supercat® Swissinno, Switzerland) strategically positioned in burrow systems and left active for 24 hours, with assistance from local farmers. These traps typically cause immediate death by head trauma. In instances where this outcome did not occur, cervical dislocation was applied swiftly, in line with the guidelines of Directive 2010/63/EU concerning the welfare of animals used for scientific purposes.

As fossorial water voles is considered one of the most serious agricultural pests in grasslands and orchards in Spain (Somoano 2020), population control is mandated under Article 15 of Spanish Law 43/2002 on plant health (BOE 2008). The actions carried out in this study fall under the category of zootechnical purposes (Real Decreto 53/2013), thereby exempting the research from requiring ethical approval. Nevertheless, all procedures adhered to the principles outlined in European Directive 2010/63/EU for the protection of animals used in scientific work. Once captured, the specimens were transferred to the Anatomy Department at the Faculty of Veterinary Medicine for necropsy, with transportation times always kept under two hours.

### Olfactory bulbs extraction

To fully extract the brain, the mandible and eyes were first detached, followed by a longitudinal incision along the dorsal surface of the skull to enable the removal of the skin and underlying muscle layers. The cranial vault and nasal bones were carefully excised using gouge forceps, exposing the cerebral hemispheres and the olfactory bulbs. The next step involved delicately isolating the olfactory bulbs from their position within the ethmoidal fossa. These laminated structures are particularly fragile, and the olfactory and vomeronasal nerves, which traverse the cribriform plate of the ethmoid bone, anchor them tightly to the dura mater. The bony walls of the fossa were carefully removed, and the olfactory nerves were severed. Following this, the optic nerves, internal carotid arteries, pituitary stalk, and cranial nerves emerging from both sides of the brainstem were severed, freeing the brain. The entire brain was then immersed in freshly prepared Bouin’s fixative. This fixative is highly recommended for neuroanatomical studies due to its excellent tissue penetration and ability to preserve the integrity of soft, delicate structures, facilitating subsequent processing (Torres et al. 2021). After 24 hours of fixation, the specimens were transferred into 70% ethanol for storage.

### Processing of samples for microscopic study

#### Paraffin embedding

The process was carried out stepwise by submerging the samples sequentially in distilled water, followed by immersion in 70% ethanol, 90% ethanol, 96% ethanol, and finally pure ethanol (three washes). The tissues were subsequently clarified using a 1:1 ethanol-xylene solution, followed by two rinses in pure xylene. After this treatment, the tissue became permeable to paraffin infiltration, which was done at 60°C for at least three hours. The sample was then encased in paraffin using a dispenser to create a solid block.

#### Sectioning

Tissue sections were prepared using a rotary microtome set to a thickness of 8 µm. For each animal, the right olfactory bulb was entirely sectioned in a transverse plane, while the left one was sliced in a sagittal plane, the latter providing a clearer and more comprehensive view of the laminar structure of the AOB.

### General histological staining

To observe the various tissue components, hematoxylin-eosin (HE) was utilized as a general stain, while the Nissl stain was applied to highlight the soma and initial segments of neuronal processes. In all the specimens, the findings were consistent in terms of structural integrity and immunohistochemical patterns.

### Immunohistochemical staining

The initial step involves inhibiting endogenous peroxidase activity to ensure that the staining is specific to the primary antibody binding site. This is accomplished by incubating the slide in a 3% H_2_O_2_ solution in distilled water, which depletes endogenous peroxidase and renders it inactive. Following three rinses in 0.1 M phosphate buffer (PB) at pH 7.2, the next step is to prevent non-specific bindings. For this purpose, a blocking serum is applied from the same species as the one used to produce the secondary antibody (ImmPRESS™ Anti-Rabbit and Anti-Mouse Kit, Vector Labs, Burlington, USA). The blocking process was carried out at room temperature for at least 20 minutes.

The immunohistochemical analysis included a specific set of antibodies that provided valuable insights into the morphofunctional characteristics of the VNS. The data for each antibody used is presented in Table 1. Antibodies against Gαi2 and Gαo were utilized to identify which pheromone receptor families—V1R (Shinohara et al. 1992) or V2R (Ortiz-Leal et al. 2024), respectively—are expressed in the VNS. Mitral cells, the primary neural components of the AOB, were marked using antibodies against microtubule-associated protein 2 (MAP-2) (Johnson and Jope 1992) and SMI-32, a marker of neurofilament proteins expressed in the mitral cells of the OB in certain therian mammals (Lee et al. 1988). Neuronal growth was assessed using antibodies against growth-associated protein 43 (GAP-43) (Verhaagen et al. 1989). The maturity of the system was evaluated with an antibody for olfactory marker protein (OMP) (Smithson and Kawaja 2009). Calcium-binding proteins, calbindin (CB) and calretinin (CR), were employed to distinguish neuronal subpopulations (Ortiz-Leal et al. 2022a). Astrocytes and ensheathing cells were identified using an antibody against glial fibrillary acidic protein (GFAP) (Taroc et al. 2020).

**Table 1.** Lectins used, concentrations and binding specificities.

| Lectin | Abbrev. | Dilut.m<br>g/ml | Supplier<br>Catalog<br>number | Preferred<br>sugar<br>specificity | Specificity<br>groups |
| --- | --- | --- | --- | --- | --- |
| *Ulex europaeus*<br>(Gorse)<br>agglutinin | UEA | 2.0 | Vector<br>B-1065-2 | α-Fuc | Fucose |
| *Lycopersicon*<br>*esculentum*<br>(Tomato) lectin | LEA | 1.0 | Vector<br>B-1175-1 | β-1,4<br>GlcNAc<br>oligomers | GlcNAc |
| *Vicia villosa*<br>(Hairy Vetch)<br>agglutinin | VVA | 2.0 | Vector<br>B1235-2 | GalNAc | Gal/GalNAc |
| *Solanum*<br>*tuberosum*<br>(Potato) lectin | STA | 2.0 | Vector<br>B-1165-2 | GlcNAc<br>Oligomers,<br>LacNAc | GlcNAc |
| *Dolichos*<br>*biflorum* (Horse<br>gram) lectin | DBA | 2.0 | Vector<br>B-1035-5 | $\alpha$ GalNAc | GalNAc |
| Abbreviations: Fuc, fucose; Gal, galactose; GalNAc, N-acetyl-galactosamine; GlcNAc, N-acetyl-glucosamine; LacNAc, N-acetyl-lactosamine. |  |  |  |  |  |

Following an overnight incubation at 4°C, the samples were rinsed three times in phosphate buffer (PB). Depending on the blocking agent applied, they were then incubated for 30 minutes with either ImmPRESS VR Polymer HRP Anti-Rabbit IgG or Anti-Mouse IgG reagents. Before proceeding to the visualization step, all samples were washed for 10 minutes in 0.2 M Tris-HCl buffer (pH 7.61). For visualization, a chromogen solution of 3,3-diaminobenzidine (DAB) was used. The reaction was developed using a 0.003% H2O2 and 0.05% DAB solution in 0.2 M Tris-HCl buffer, producing a brown precipitate. Negative controls were prepared by omitting the primary antibodies.

### Lectin histochemical labelling

Lectin labeling, while similar to immunohistochemical techniques, relies on specific proteins known as lectins. These proteins have domains that recognize and bind non-covalently to terminal sugars in tissues, forming glycoconjugates. Unlike antibodies, lectins do not have an immune origin (Lis and Sharon 1998). Lectins have been extensively used in the study of the olfactory bulb (Shin et al. 2017). In this study, we employed five lectins (Table 1).

#### Ulex europaeus agglutinin

UEA specifically binds to terminal L-fucose present in glycoproteins and glycolipids. It was chosen for this study because of its role as a specific marker for the AOB in several species, including mice (Kondoh et al. 2017), fox (Ortiz-Leal et al. 2022b), and wolves (Ortiz-Leal et al. 2024).

#### Lycopersicum esculentum agglutinin

LEA exhibits high affinity for N-acetylglucosamine and has been widely used as a comprehensive marker for both the main and accessory olfactory bulb in various species. These include rodents such as rats (Franceschini et al. 1994), mice (Keller et al. 2022), capybaras (Torres et al. 2020), and hamster (Taniguchi et al. 1993b).

#### Vicia villosa agglutinin

VVA, which is isolated from fodder vetch, binds specifically to N-acetylgalactosamine structures (Tomiyasu et al. 2018). In rats, it selectively stains the accessory olfactory bulb (Ichikawa et al. 1992; Takami et al. 1992), but this is not the case in mice (Salazar and Sánchez Quinteiro 2003).

#### Solanum tuberosum agglutinin

STA, which is isolated from potato, binds oligomers of N-acetylglucosamine and some oligosaccharides containing N-acetylglucosamine and N-acetylmuramic acid (Chun et al. 2023).

#### Dolichos biflorus agglutinin

DBA is isolated from horse gram and has a carbohydrate specificity toward N-acetylgalactosamine (Torres et al. 2023).

The lectin histochemical labelling process began with slides that had been deparaffinized and rehydrated. To suppress endogenous peroxidase activity, they were treated with a 3% H2O2 solution. A 2% bovine serum albumin (BSA) solution was applied to block non-specific bindings. The lectins were incubated overnight at 4°C. Following this, the slides were exposed to a 90-minute incubation in an avidin-biotin complex (ABC) solution with peroxidase (Vectorlabs, Burlingame, CA). This complex interacts with the lectin during the incubation, enhancing the subsequent peroxidase reaction. The reaction was visualized using a solution of 0.003% H2O2 and 0.05% 3,3-diaminobenzidine (DAB) in 0.2 M Tris-HCl buffer, which produced a brown precipitate. Controls included tests without the lectin, as well as with lectins pre-absorbed with an excess of their corresponding sugars

### Image acquisition

Digital images were acquired using an Olympus SC180 camera attached to an Olympus BX50 microscope. To create composite images from multiple photographs, PTGuiPro software (Rotterdam, Netherlands) was employed due to the large size of the studied structures. Adobe Photoshop CS4 (San Jose, CA) was utilized to modify brightness, contrast, and white balance levels; however, no enhancements, additions, or alterations to the image characteristics were performed.

## Results

The macroscopic study of the brain was conducted after its complete extraction (Fig. 1), which was performed following the removal of the skin of the head and the dorsal opening of the cranial vault. The main olfactory bulbs exhibit a development comparable to that of more extensively studied rodents, such as rats and mice. They are located at the rostral part of the brain, positioned anteriorly to the other structures, and are clearly visible as two projections extending from the telencephalon (Fig. 1A-C). The AOB is not easily recognizable macroscopically, as it does not form a prominent protrusion on the surface of the MOB and is also concealed by the frontal lobe of the telencephalon. Therefore, the best way to visualize it is by ventrally pulling the MOB with the aid of a forceps. The ventral view of the brain allows for the observation of the significant development of the basal part of the rhinencephalon, which is associated with the olfactory subsystems (Fig. 1C). It is remarkable the presence of a wide band of the lateral olfactory tract connecting the OB with the piriform lobe.

**Figure 1.**
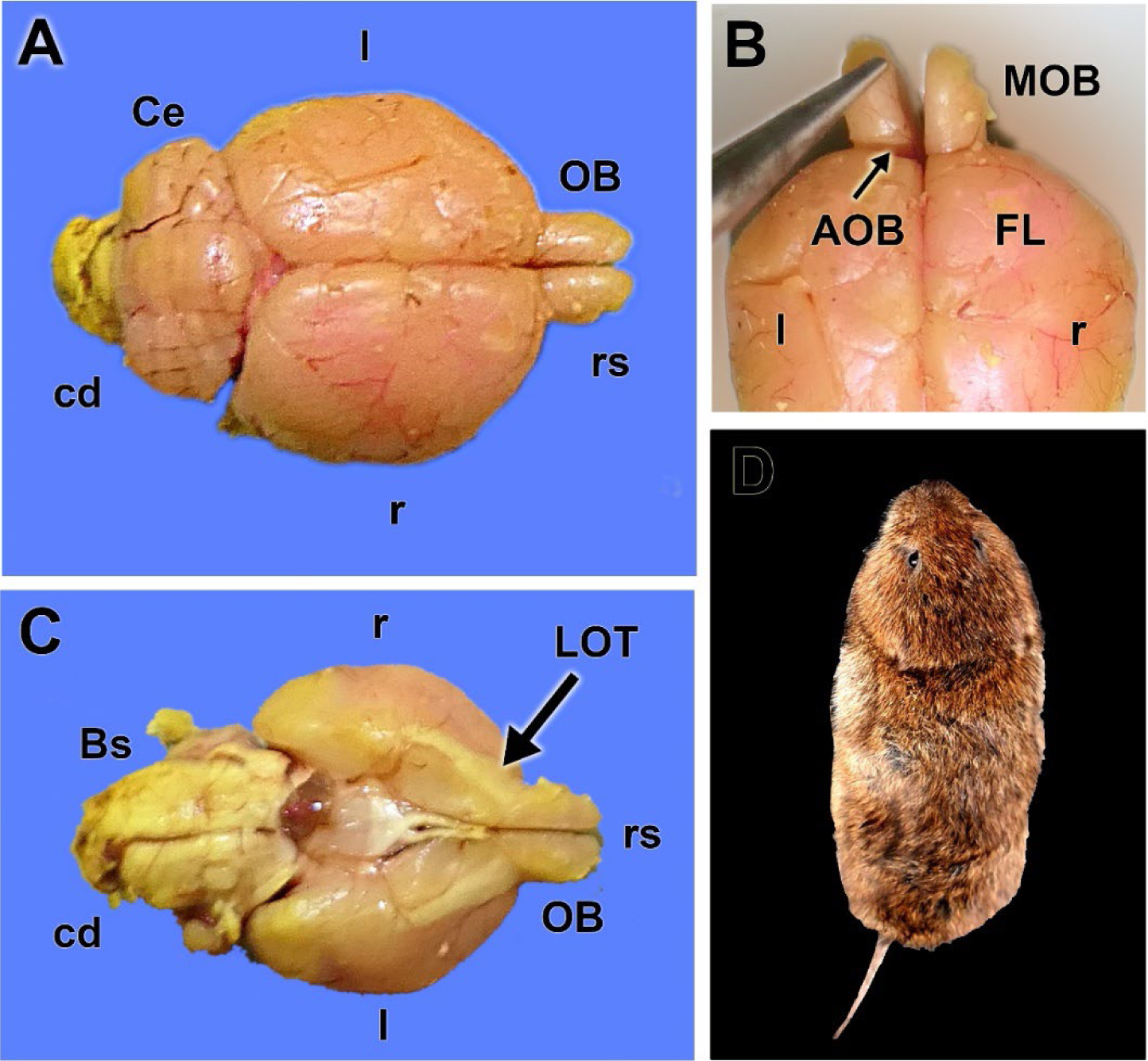
Macroscopic anatomy of the olfactory bulb of the fossorial water vole. **A.** Dorsal view of the brain showing the relative size of the olfactory bulbs (OB). **B.** Dorsal view of the rostral part of the telencephalon showing the location area of the AOB. **C.** Ventral view of the brain showing the significant development of the lateral olfactory tract (LOT). **D.** Dorsal view of one of the analyzed specimens of fossorial water vole. Bs: Brainstem, cd: Caudal, Ce: Cerebelum, FL: frontal lobe, l: left, MOB: main olfactory bulb, r: right, rs: Rostral.

The histological study enabled the identification and exhaustive characterization of the accessory olfactory bulb of the fossorial water vole. It presents a well-defined laminar structure, similar to the main olfactory bulb, although there are remarkable differences between them, as shown in Fig. 2A, B. The nervous Layer (NL) of both structures receives vomeronasal and olfactory axons, respectively for the AOB and MOB. The glomerular structures are clearly defined in both structures. They are the destination of the sensory axons, and the synapsis point with dendrites of the subsequent neuron in the olfactory and vomeronasal pathway, the mitral cells. However, the MOB shows a much higher density of periglomerular cells (PGC).

**Figure 2.**
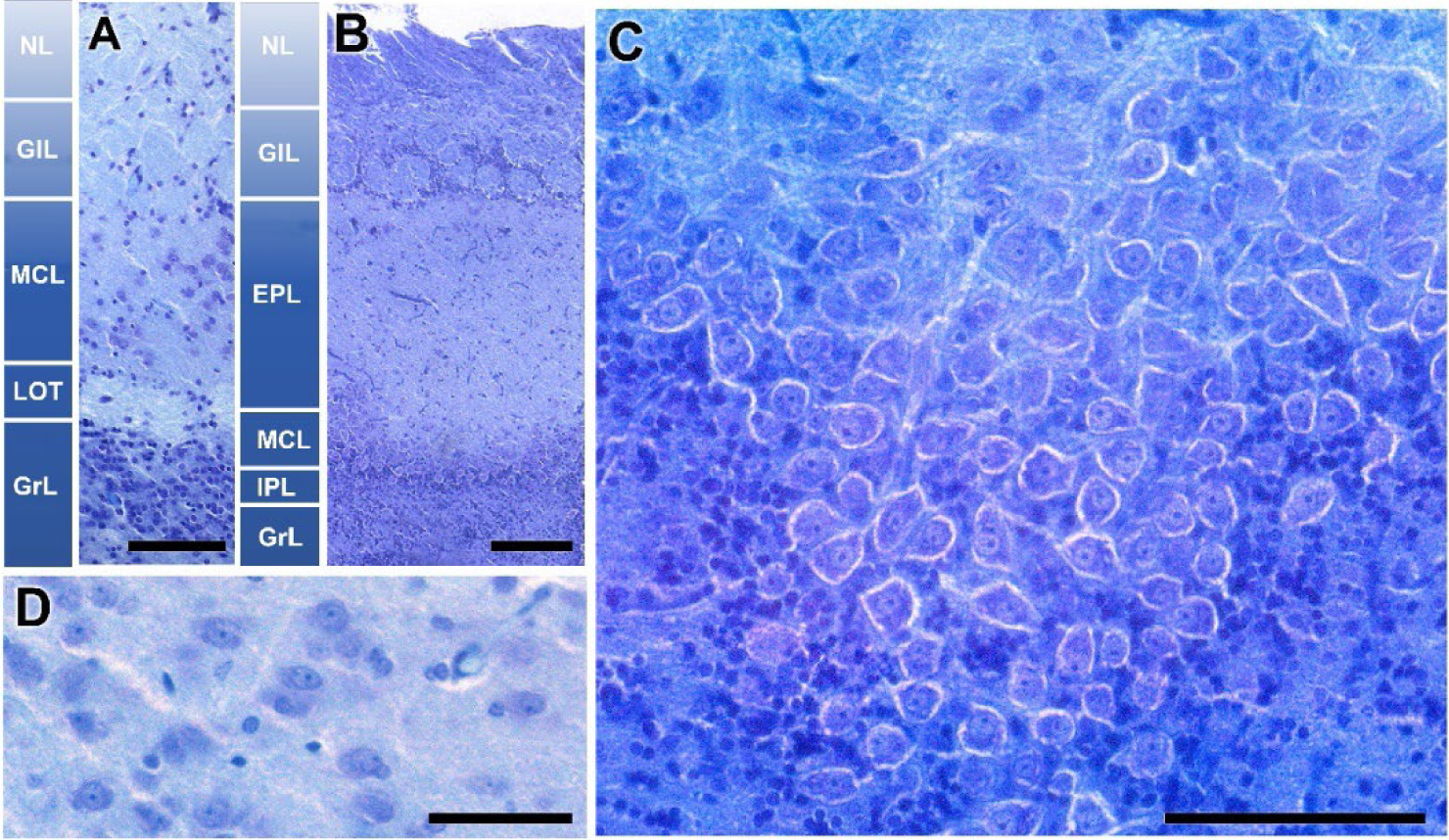
Histological structure of the olfactory bulb of the fossorial water vole. **A-B.** Comparative study of the lamination of the accessory olfactory bulb and the main olfactory bulb, respectively. **C.** Horizontal section of the mitral cell layer of the MOB. **D.** Mitral cells of the AOB. Nissl Staining. EPL: Externa plexiform layer, GlL: Glomerular layer, GrL: Granular layer, IPL: Plexiform internal layer, LOT: Lateral olfactory tract, MCL: Mitral cells layer, NC: Nervous layer. Scale bars: A, B, C = 100 µm, D = 50 µm.

The most remarkable difference is observed in the intermediate layers of the bulb, which contain the mitral cells. In the MOB, these are arranged along an axis forming an individualized lamina (MCL). In a horizontal section of this layer in the MOB, these cells exhibit significant development, with an atypical configuration not seen in other mammalian species (Fig. 2C). Above this layer, there is a wide neuropil made up of the MC dendrites, known as the external plexiform layer (EPL). Its deeper counterpart is a thin layer, the internal plexiform layer (IPL), containing the beginnings of the axons of these mitral cells. However, the organization of the second-order neurons in the vomeronasal pathway is very different. The neuronal somas are not mitral or polygonal in shape, but oval (Fig. 2D), and they are not arranged in a continuous lamina but are dispersed, equivalent to the EPL of the MOB. Therefore, we adopt the term mitral-plexiform layer (MPL) to design the resulting layer from the fusion of the two plexiform layers and the mitral layer. Deeper to the mitral plexiform layer is a specific layer of the BOA; the one that forms the lateral olfactory tract (LOT). This consists of a cord of white matter formed by the axons of mitral cells from both the MOB and the AOB. In the deepest plane, a broad layer of granular cells (GrL) is present in both systems.

The AOB was examined through Nissl-stained sagittal serial sections, to comprehensively characterize its laminar organization (Fig, 3). The correlative sections illustrate from medial to lateral the structural organization of the AOB (Fig. 3A-G). The glomerular, mitral-plexiform, and granular layers are clearly distinguishable, as outlined by a yellow dashed line in Fig. 3D.

**Figure 3.**
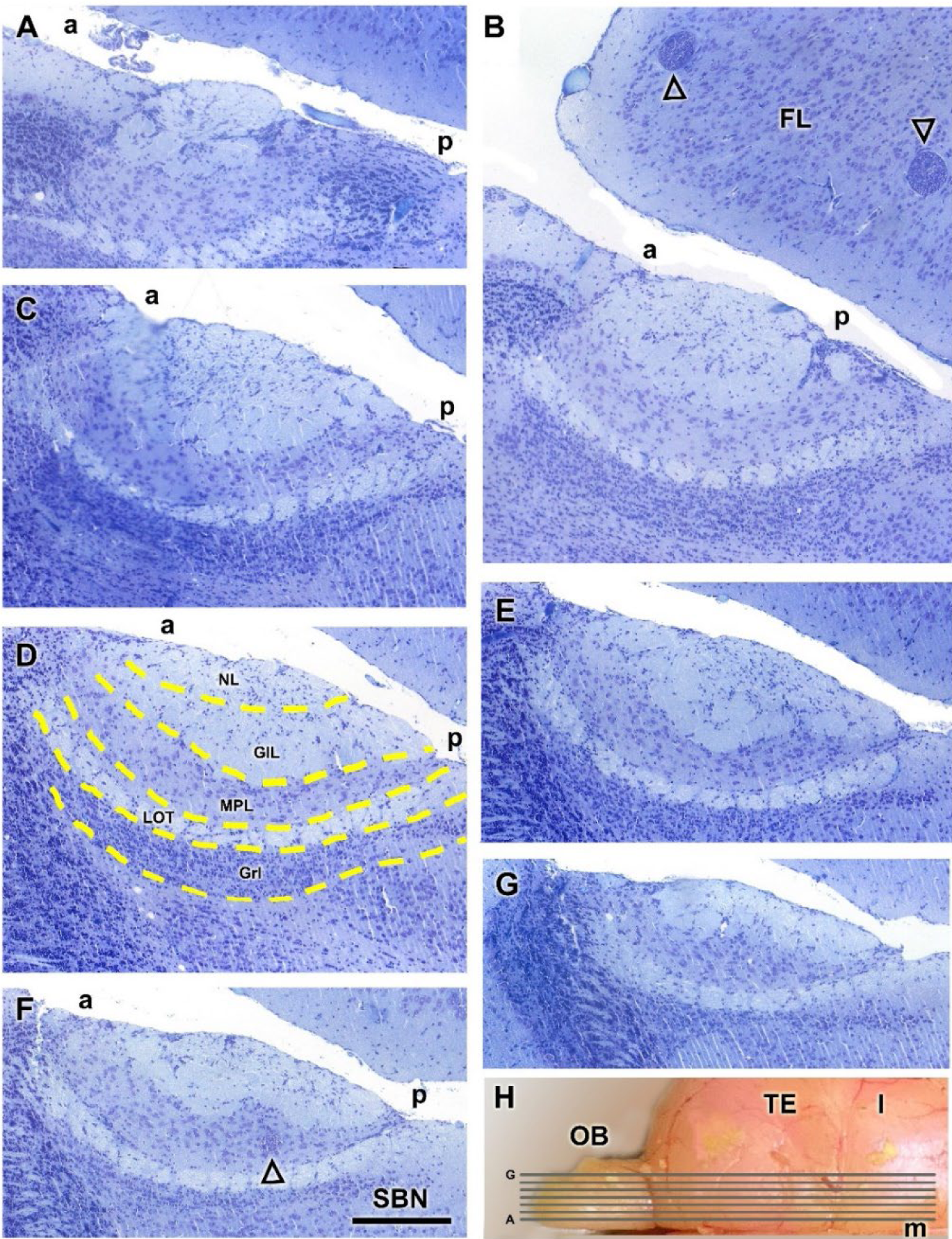
Nissl staining of sagital serial sections of the accessory olfactory bulb of the fossorial water vole. **A-G**. Correlative sections of the AOB arranged from medial to lateral. In **D**, the layers of the AOB are outlined by a yellow dashed line. In **B,** the open arrowhead represents parasitic cysts. **H**. Dorsal view of the right hemisphere of the water vole, showing the corresponding levels of sections A-G. a, anterior; FL, frontal lobe; GlL; glomerular layer; GrL, granular layer; MPL, mitral plexiform layer; l, lateral; LOT, lateral olfactory tract; m, medial; NL, nerve layer; OB, olfactory bulb; p, posterior; TE, telencephalon. Scale bar = 250 µm.

Remarkably, the glomerular and mitral-plexiform layers exhibit a significant development. Incidentally, parasitic cysts, presumably caused by *Toxoplasma* spp., were identified in the frontal lobe (Fig. 3B) and the MPL of the AOB (Fig. 3E). The presence of such cysts is consistent with prior reports of parasitic infections in wild rodents. The lateral olfactory tract is visualized all through the AOB (Fig. 3B-G).

### Lectin histochemical staining

Lectin staining of the AOB and MOB using UEA and LEA revealed distinct labeling patterns across different layers and regions. UEA staining enabled the identification of the superficial nerve and glomerular layers of the AOB (Fig. 4A-D). A clear rostro-caudal zonation was observed in the AOB, with the anterior portion showing strong positive staining, while the posterior portion remained negative (Fig. 4A). However, this zonation was not consistent across all sagittal planes of the AOB, as demonstrated in Fig. 4B, where the anterior-posterior zonation is not visible. However, a hematoxylin counterstained section shows how some isolated AOB glomeruli remain unstained by the lectin (Fig. 4C). In addition to variability in AOB staining, differences were observed in the staining pattern of the MOB depending on the specimen. Some individuals displayed no UEA staining in the MOB (Fig. 4A, B), whereas others exhibited a strongly positive staining pattern (Fig. 4D).

**Figure 4.**
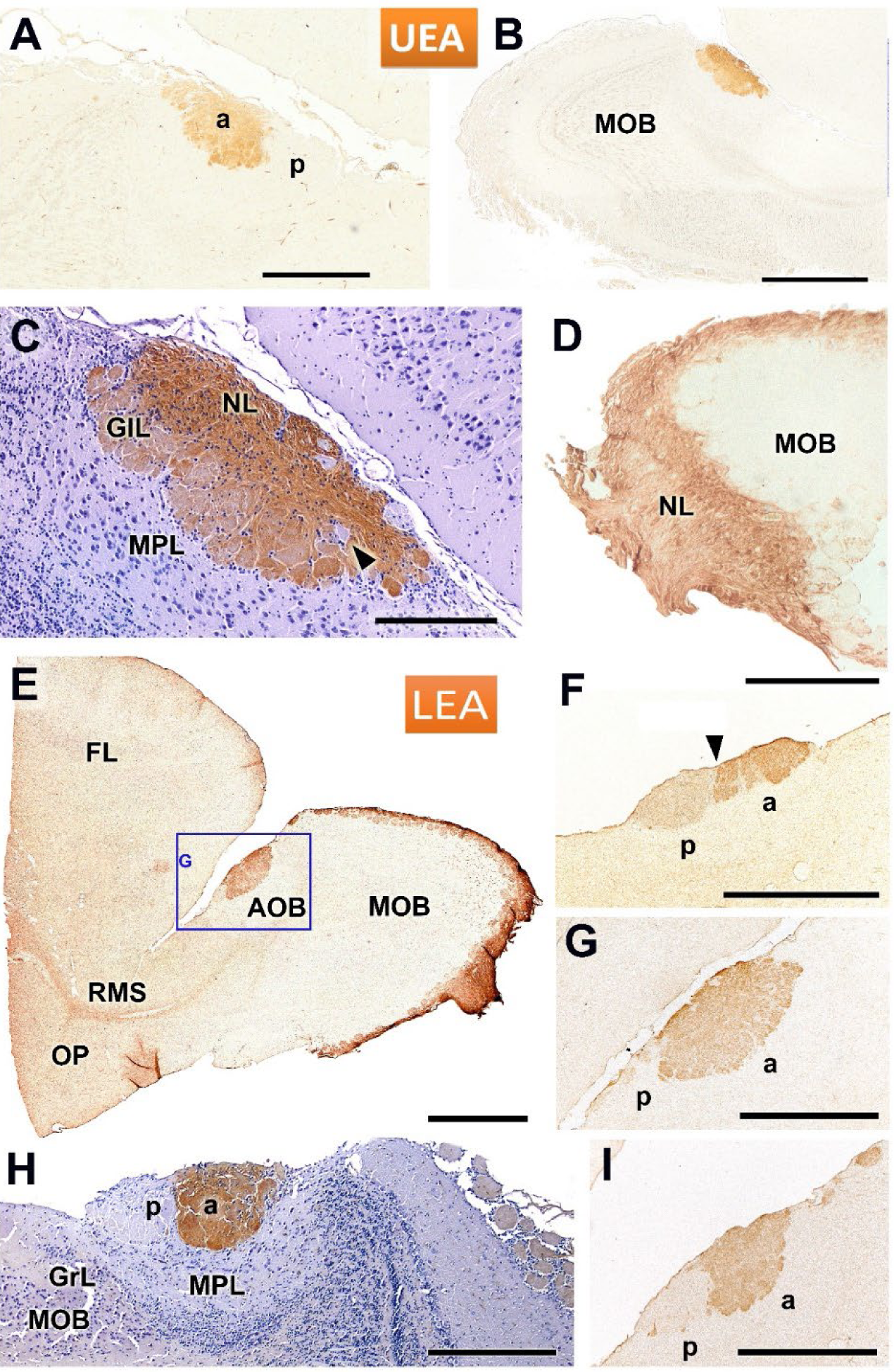
Lectin staining of the accessory and main olfactory bulbs in the fossorial water vol using UEA and LEA. **A-D**. UEA staining allows for the identification of the nerve and glomerular layers of the AOB. **A.** Sagital section shows the presence of a rostro-caudal zonation in the AOB, with a positive anterior portion and a negative posterior portion. This zonation does not extend to all sagital planes of the AOB, as shown in images **B** and **C**. A hematoxylin counterstained section (C) shows how some isolated AOB glomeruli remain unstained by the lectin (arrowhead). Depending on the specimen studied, there are significant differences in the staining patern of the MOB. Some individuals are UEA-negative for the MOB (A, B) while others exhibit a positive patern (D). **E-I**. LEA staining consistently showed a positive patern in the superficial layers of both the AOB and MOB. **E**. Overview of the anterior telencephalon, showing strong LEA staining in the AOB (highlighted in **G**) and the MOB, along with the rostral migratory stream (RMS). **F, G, I**. Sagital sections of the same AOB at different levels reveal an anterior-posterior zonation (arrowhead in E). The intensity of the zonation depends on the level considered. **H**. Counterstained image from a different individual confirming the zonation and showing how the staining encompasses the glomerular (GlL) and nerve (NL) layers of the anterior AOB. a, anterior; FL, frontal lobe; p, posterior. Scale bars: B, E = 1 mm; A, D-G, I = 500 µm; C, H = 250 µm.

LEA staining consistently revealed a positive labeling pattern in the superficial layers of both the AOB and MOB in all cases examined. An overview of the anterior telencephalon showed robust LEA staining in the superficial layers of both the AOB and MOB, as well as in the rostral migratory stream (RMS) (Fig. 4E). Sagittal sections of the AOB at different rostro-caudal levels revealed an anterior-posterior zonation in the LEA staining, although the degree of this zonation varied depending on the specific level observed (Fig. 4F, G, I). A counterstained image from a different individual confirmed the presence of this zonation, with staining encompassing both the glomerular (GlL) and nerve (NL) layers in the anterior AOB (Fig. 4H).

The distribution of lectin binding in the olfactory bulb (OB) with lectins DBA, STA, and VVA is shown in Fig. 5. DBA staining was localized exclusively to the AOB, demonstrating a clear delineation between the AOB and the MOB, which remained unstained (Fig. 5A). STA lectin resulted in a staining pattern that resembled that observed with LEA, prominently staining the superficial layers of both olfactory bulbs (Fig. 5B-D). The figures shown correspond to two individuals, one of them counterstained with hematoxylin (Fig. 5C). Both specimens confirmed the presence of anteroposterior zonation within the AOB (Fig. 5C, D). Finally, VVA lectin exhibited generalized nuclear staining within the OB, in the AOB predominantly concentrated in its mitral plexiform and granular layers (Fig. 5E).

**Figure 5.**
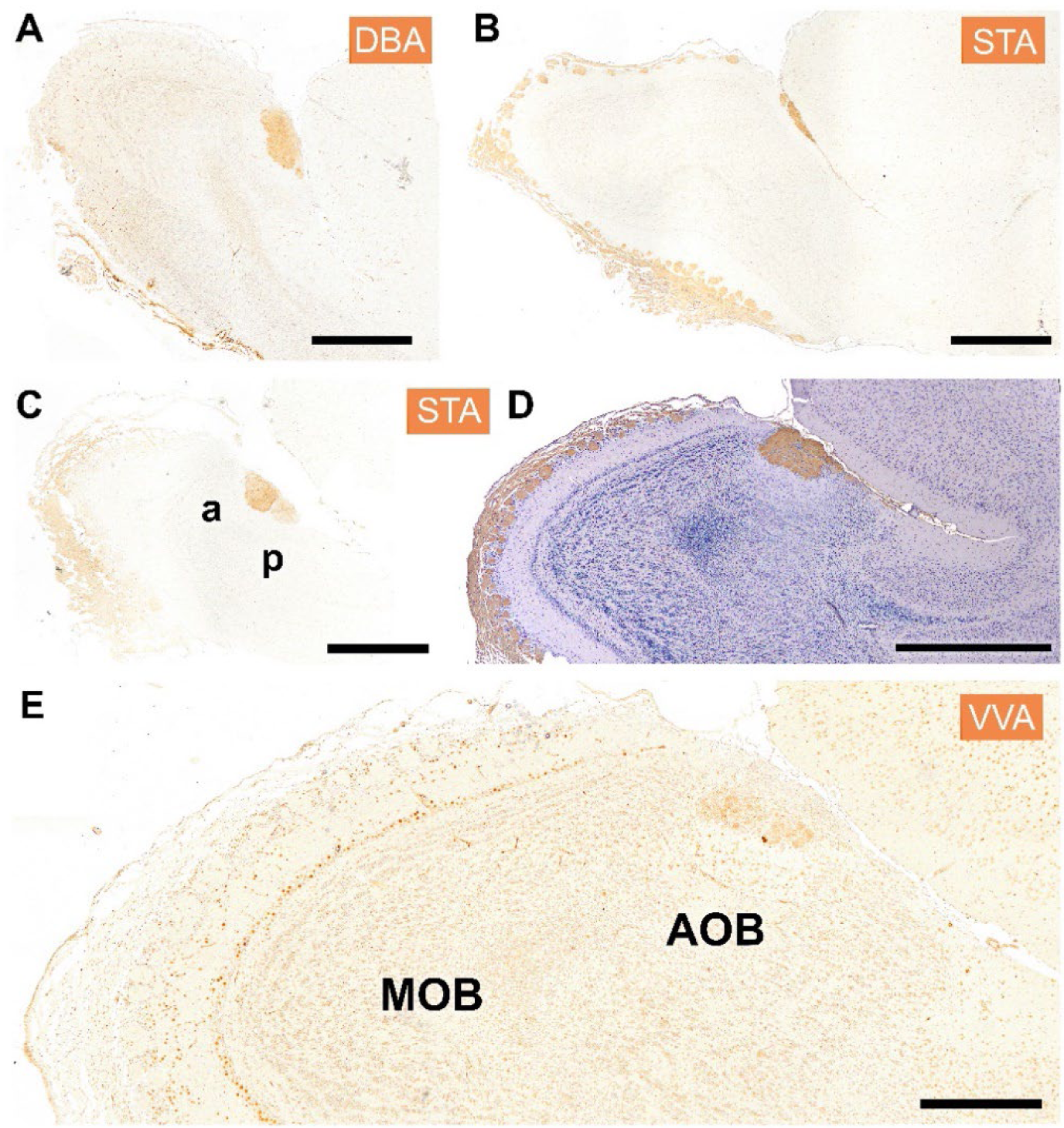
Histochemical staining of the fossorial water vole OB with the lectins DBA, STA, and VVA. **A.** DBA specifically stains the whole AOB as it does not stain the MOB. **B-D.** STA shows a similar staining patern to that observed with the lectin LEA, with strong staining in the superficial layers of both the main and accessory OB. B and D correspond to two sagital levels of the same individual, and C to a second specimen. Both individuals show anteroposterior zonation (C, D). E. The lectin VVA shows generalized nuclear staining, mostly concentrated in the mitral plexiform and granular layer but involving a similar patern to the MOB. Scale bar: A-D = 1 mm; E = 500 µm.

### Immunohistochemical study

The immunohistochemical study allowed for a deeper understanding of the morphofunctional characterization of the VNO. The study of the protein subunits of G protein, Gα0 and Gαi2, is crucial due to their direct relationship with the expression of the vomeronasal receptors of the V1R and V2R families. Gα0 proteins are primarily associated with the V2R family, while Gαi2 proteins are related to the V1R family. In the AOB of the fossorial water vole, immunolabeling with anti-Gαo antibody shows a strong immunostaining in the posterior zone of the AOB, as well as in the superficial layers of the MOB (Fig. 6A). The enlarged view of the AOB shows a clear demarcation from anterior and posterior regions of the AOB at the level of the superficial layers, nervous and glomerular (Fig. 6B).

**Figure 6.**
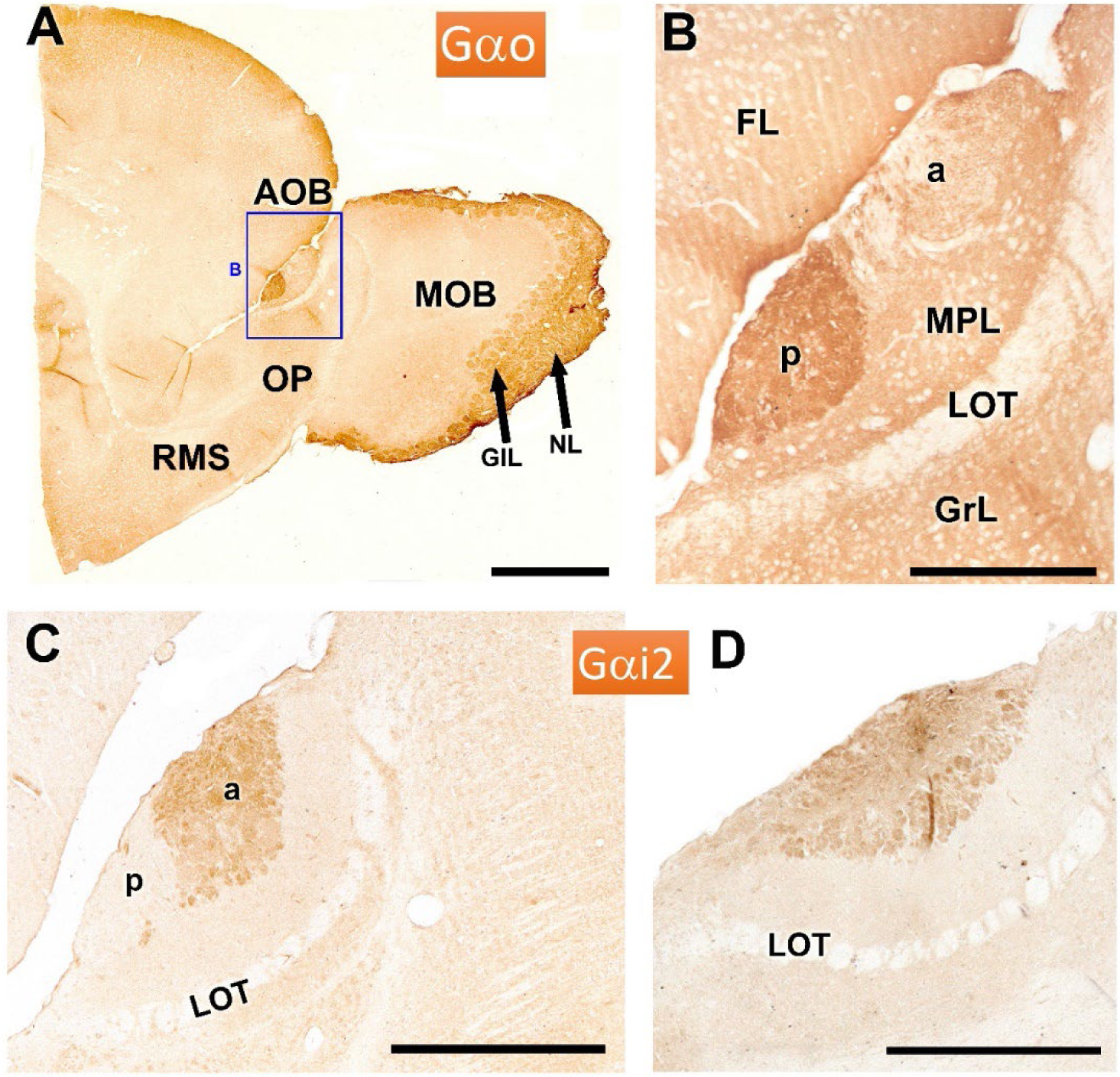
Immunohistochemical labeling of the AOB of the fossorial water vole with antibodies against the G protein subunits αo and αi2. **A**-**B**. Immunolabeling with anti-Gαo. The anterior part of the telencephalon shows intense immunopositivity in the posterior zone of the AOB (enlarged in B) and in the superficial layers of the MOB. **C-D**. Immunolabeling with anti-Gαi2 in two sagital sections of the AOB shows anteroposterior zonation only in one of them (C), with the immunolabeling concentrated in the anterior part of the AOB. The more lateral section (D), however, does not show such zonation. a: anterior, GlL: Glomerular layer, GrL: Granular layer, LOT: lateral olfactory tract, MPL: Mitral plexiform layer, NL: Nervous layer, p: posterior, OP: Olfactory peduncle, RMS: rostral migratory stream. Scale bars: A = 1 mm, C = 500 µm, B, D = 250 µm.

Immunolabeling with anti-Gαi2 reveals a complementary anteroposterior zonation to that observed with anti-Gαo, with the immunolabeling in this case concentrated in the anterior part of the AOB (Fig. 6C). In contrast, a more lateral section does not exhibit such zonation, highlighting variability in the expression across the sagittal plane of the AOB (Fig. 6D).

Immunohistochemical analysis using antibodies against calbindin (CB) and calretinin (CR) revealed distinct patterns of labeling in the AOB and MOB (Fig. 7). Anti-CB staining produced a strong labelling in the nerve (NL) and glomerular (GL) layers of the AOB. However, the mitral plexiform (MPL) and granular (GrL) layers also displayed positivity, albeit with a weaker intensity (Fig. 7A). Additionally, a nucleus of strong immunopositive neurons was detected in the subbulbar region (SBN) (Fig. 7B). In the MOB, CB immunostaining was particularly intense in the nerve layer, but the glomerular neuropil remained unstained. Nevertheless, periglomerular cells showed strong immunopositivity (Fig. 7C).

**Figure 7.**
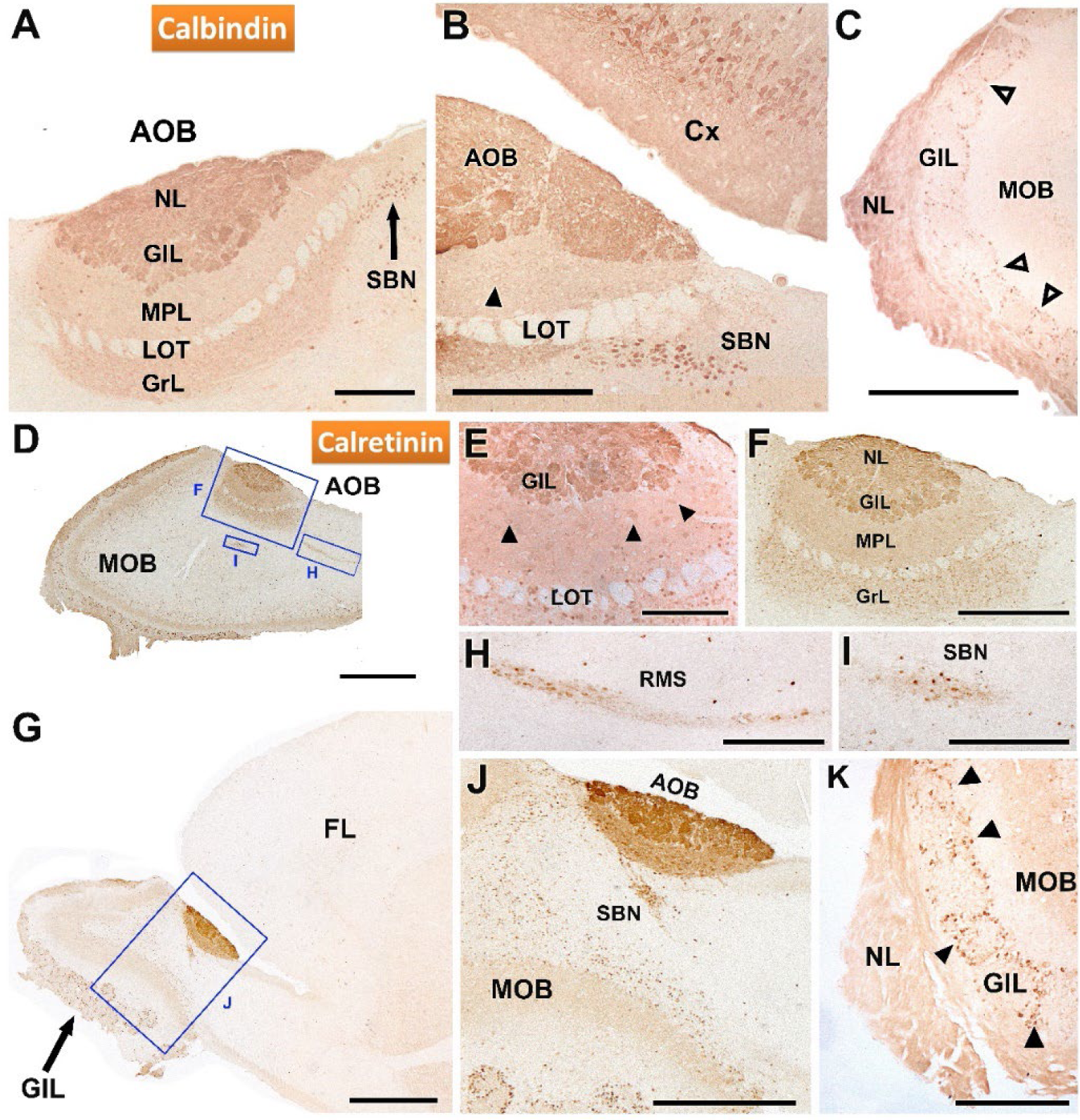
Immunohistochemical labeling of the AOB of the fossorial water vole with antibodies against the calcium binding protein calbindin (CB) and calretinin (CR). **A-C.** Immunostaining with anti-CB. **A.** Immunostaining in the AOB is concentrated in the nerve and glomerular layers, although the mitral plexiform and granular layers are also positive but with a weaker signal. Additionally, a nucleus of immunopositive neurons is identified in a subbulbar position (SBN). **B.** Enlargement of the caudal area of the AOB shown in A. The neuronal somas of the SBN and the pyramidal neurons of the cerebral cortex (Cx) show strong immunopositivity. The mitral cells of the AOB (arrowheads) are weakly stained. **C.** The MOB shows intense staining in the nerve layer, does not stain the glomeruli, but does mark the periglomerular cells (open arrowheads). **D-K** Anti-CR immunostaining. **D.** Sagital section of the OB showing immunopositivity in the AOB and MOB. **E.** Anti-CR produces intense staining in the nerve and glomerular layers, and in the mitral cells of the AOB (arrowhead). **F.** Enlargement of the inset in C showing immunostaining comprising all layers of the AOB, being especially intense in the two superficial layers, nerve (NL) and glomerular (GL). **G.** Low magnification sagital view of the immunostaining in the anterior part of the telencephalon. **H-I**. Enlargements of the boxes in G showing, respectively, the intense immunopositivity in the RMS and the SBN. **J.** Enlargement of the box in G showing the direct connection between the AOB and the RMS. **K.** Immunostaining in the MOB extends to the nerve layer and the periglomerular cells (arrowhead). GcL: Granular layer, MPL, mitral-plexiform layer; RMS: Rostral migratory stream, LOT: Lateral olfactory tract. Scale bars: 1 mm = D, G, 500 µm = J, 250 µm = A-C, F, H, I, 50 µm = E.

Immunohistochemistry with anti-calretinin further clarified the labeling patterns in both the AOB and MOB (Fig.7D-K). Both structures showed strong immunopositivity, with strong labelling in nerve, glomerular, and granular layers of the AOB (Fig. 7D-G). Mitral cells of the AOB were also immunolabelled with anti-CR (Fig. 7E). Examination of the AOB at higher magnification revealed that immunostaining extended to all layers, though it was particularly intense in the superficial layers (NL and GL) (Fig. 7E, F). A low magnification sagittal view of the anterior telencephalon (Fig. 7G) revealed calretinin immunoreactivity in the rostral migratory stream (Fig. 7H) and in the SBN (Fig. 7I). Remarkably, a direct topographical relationship between the AOB and the RMS was evident (Fig. 7J). In the MOB, calretinin staining was prominent in the nerve layer and in periglomerular cells (Fig. 7K).

Immunohistochemical analyses using antibodies against MAP2 and GAP43 are depicted in Fig. 8A, B. Anti-MAP2, produced strong immunoreactivity in the external plexiform layer of the MOB and in the mitral-plexiform layer of the AOB, as well as in the olfactory peduncle (Fig. 8A) and the frontal lobe (Fig. 8B). Remarkably, the rostral migratory stream (Fig. 8A) and the lateral olfactory tract (Fig. 8b) exhibited no staining, indicating a lack of MAP2 expression in this area (Fig. 8A). Within the AOB, the dendrites of MPL and GrL displayed strong immunoreactivity, whereas the cellular somas remained unstained, suggesting differential localization of MAP2 within neuronal components.

**Figure 8.**
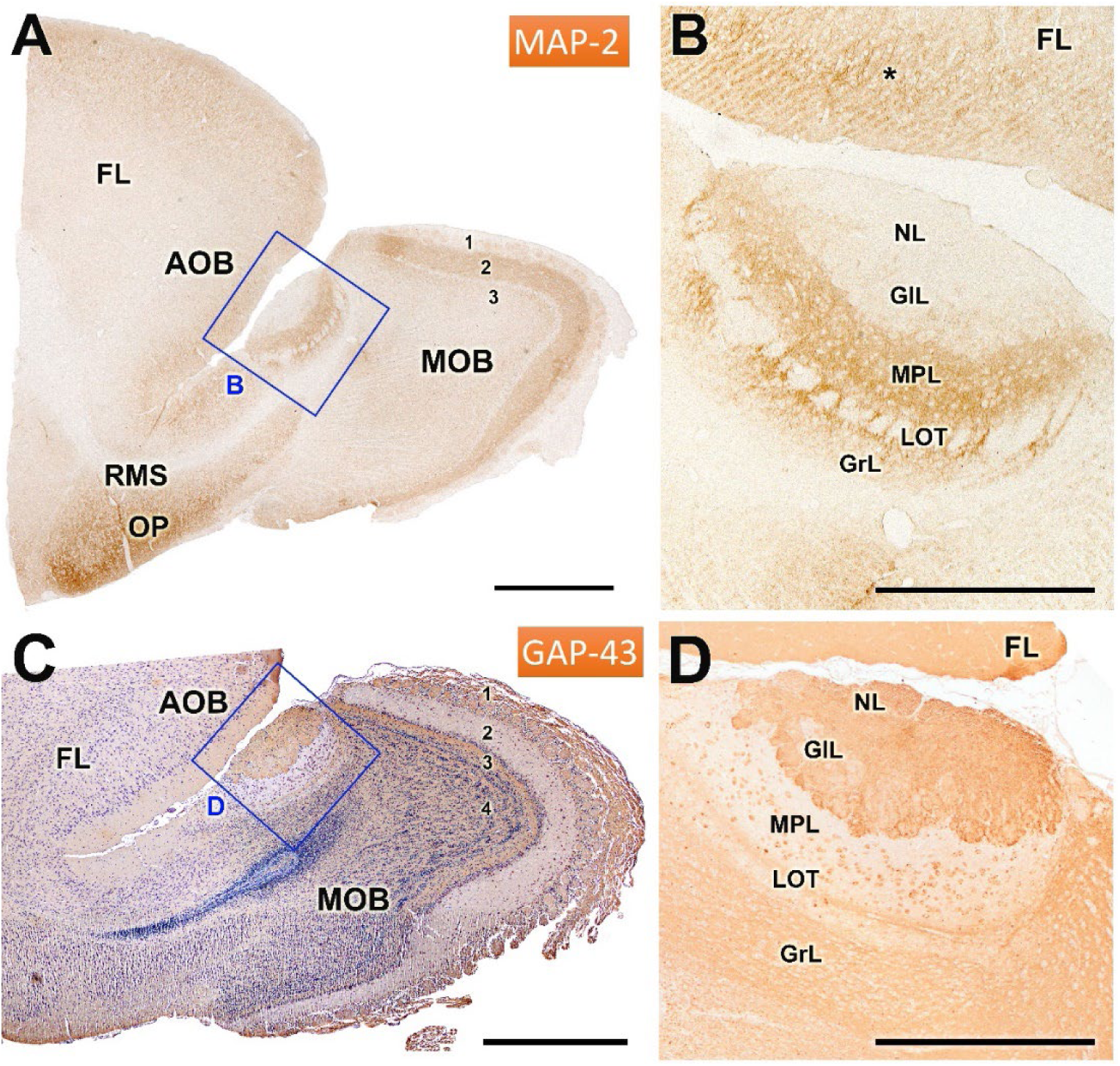
Immunohistochemical labeling of the olfactory bulb of the mole rat with anti-MAP2 and anti-GAP43. **A-B**. Immunostaining of the olfactory bulb with anti-MAP2. **A.** Sagital section of the anterior part of the telencephalon showing strong immunostaining in the external plexiform layer of the MOB (2) and the mitral-plexiform layer (MPL) of the AOB, and in the olfactory peduncle (OP). The rostral migratory stream (RMS) is immunonegative. **B**. Enlargement of the inset in A showing immunostaining in both the AOB and the frontal lobe (*). The dendrites of the mitral-plexiform (MPL) and granular layer (GrL) are strongly stained, however, the cellular somas are immunonegative. **C-D**. Immunostaining of the OB with anti-GAP43. **C**. H-E counterstained sagital section of the rostral telencephalon. The patern of anti-GAP43 immunostaining is complementary to that observed with anti-MAP2. Immunopositivity is thus observed in the superficial layers of the AOB and MOB (NL and GlL) and in the GrL of AOB and MOB. **D**. Higher magnifications of the inset in C shows strong immunopositivity in the superficial layers, and additionally, the mitral cells of the MPL layer and the lateral olfactory tract (LOT) traversing it are also immunopositive. FL: Frontal lobe, 1: Nerve + glomerular layer, 2: External plexiform layer, 3: Internal plexiform layer, 4: Granular layer. Scale bar: A, C = 1 mm; B, D = 500 µm.

The staining pattern with anti-GAP43 (Fig. 8C, D) showed a complementary pattern to anti-MAP2 immunostaining. Thus, immunopositivity was primarily concentrated in the nerve, glomerular, and granular layers of both the AOB and MOB (Fig. 8C). Additionally, in the AOB, the mitral cells within the MPL layer and the lateral olfactory tract traversing them also displayed strong immunopositivity for GAP43.

We completed the immunohistochemical analysis of the OB with antibodies against OMP, DCX, SMI32, PGP, and SG (Fig. 9). Anti-olfactory-marker-protein (Fig. 9A, B) produced a homogeneous labelling of the superficial layers of both the AOB and MOB. Thus, there was no evidence of anterior-posterior zonation. The intensity of the immunostaining was higher in the AOB than in the MOB. Anti-DCX demonstrated the presence of doublecortin in the cells belonging to the rostral migratory stream, which suggests active neurogenesis and a dynamic movement of new neurons from the subventricular zone towards the OB (Fig. 9C). In contrast, anti-SMI32 immunostaining was localized specifically to the MOB, with intense immunoreactivity observed in the somas and dendrites of neurons within the mitral and the external plexiform layer (Fig. 9D).

**Figure 9.**
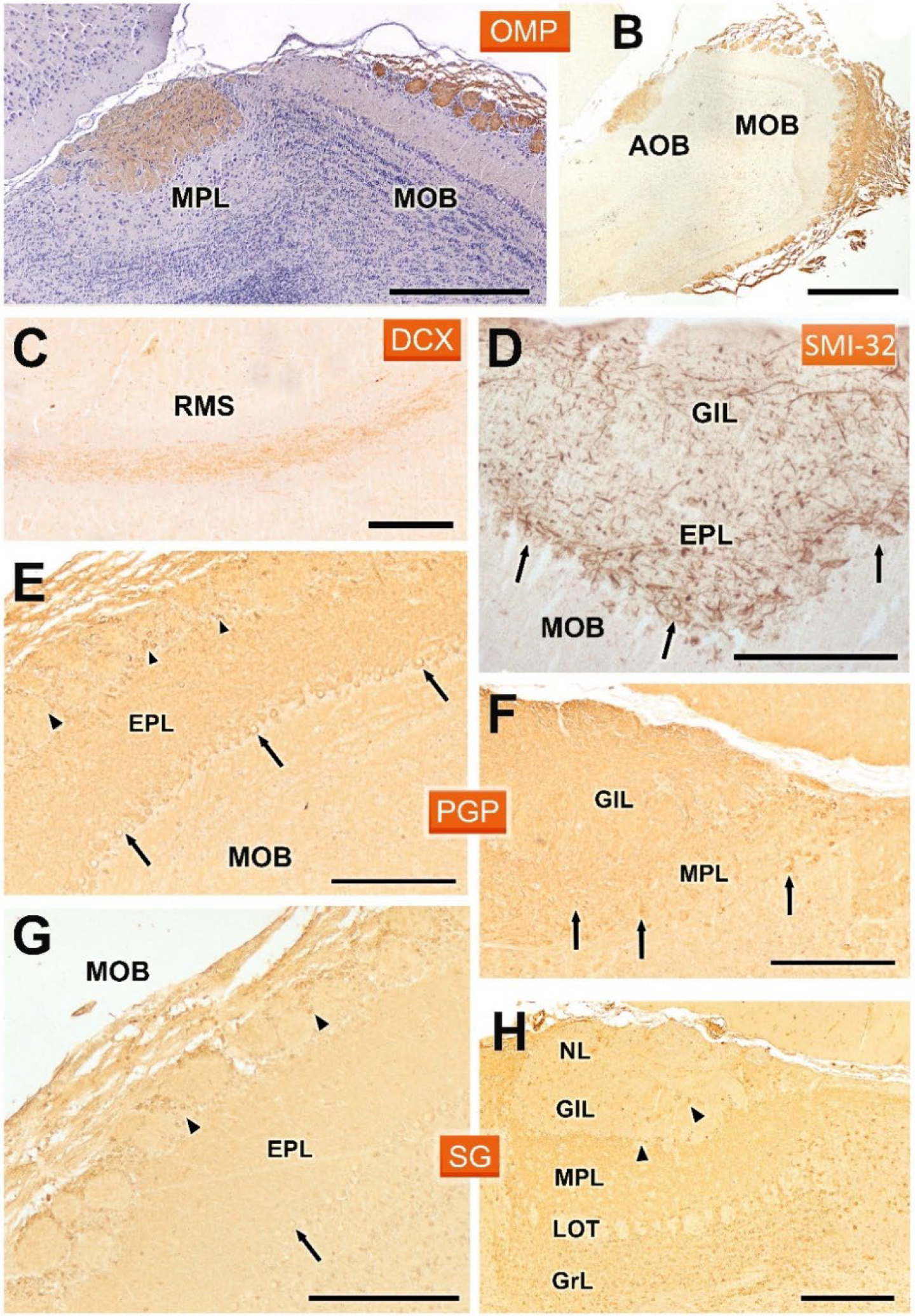
Immunohistochemical labelling of the fossorial water vole OB with antibodies against OMP, DCX, SMI32, PGP, and SG. **A-B.** Anti-OMP produces intense labeling in the superficial layers of the AOB and MOB. The hematoxylin counterstained image (A) shows the absence of anterior-posterior zonation. **C.** Anti-DCX labels the cells of the RMS. **D.** Anti-SMI32 only stains the MOB, where it intensely labels the somas (arrows) and dendrites in the mitral and external plexiform layer (EPL). **E-F.** Anti-PGP produces generalized labelling of the neuropil in both the MOB and AOB. At the cellular level, in the MOB (E) it labels the mitral (arrows) and periglomerular cells (arrowheads), whereas in the AOB (F) it produces a similar patern to that observed in the MOB, but without labelling the periglomerular cells. **G-H.** Anti-SG produces a similar patern in both structures, MOB (G) and AOB (H), although it does not stain mitral cells in the later. GlL: Glomerular layer, GrL: Granular layer, LOT: Lateral olfactory tract, MPL: Mitral-plexiform layer, NL: Nerve layer. Scale bars: 1 mm = B, 500 µm = A, 250 µm = C-H.

The staining with anti-PGP (Fig. 9E, F) and anti-SG (Fig. 9G, H) showed very similar patterns in both the MOB and AOB. In both cases, there was generalized labeling of the neuropil, reflecting widespread neuronal activity in these regions. In the MOB, anti-PGP strongly labeled mitral cells and periglomerular cells (Fig. 9E), while in the AOB a similar pattern was observed, though the periglomerular cells were not stained (Fig. 9F). Similarly, anti-SG also labeled the neuropil in both structures, although mitral cells in the AOB did not show immunoreactivity, which may suggest differences in synaptic organization and function between the MOB and AOB (Fig. 9H).

## DISCUSSION

One of the major limitations in the study of chemical communication in mammals is the scarce information available about their chemosensory systems, particularly the vomeronasal systems. This challenge is further complicated by the high morphological variability of the vomeronasal system among different mammalian groups, which makes extrapolation of findings difficult (Brennan 2001). If the available data on the structure, and especially the neurochemistry, of the VNO is limited, the situation becomes even more challenging when addressing the AOB (Ennis and Holy 2015). In rodents, for example, most studies are restricted to laboratory rodents, with only a limited number focusing on other rodent groups and, in few instances, addressing not only the classical morphological characteristics but also neurochemical expression, and the role of glycoconjugates identified through lectin binding.

For this reason, we chose to focus our attention on the AOB of the fossorial water vole, a subterranean rodent species belonging to the family Cricetidae, which probably relies heavily on its chemical senses for survival (Airoldi 1976). This study aims to bridge the gaps in our understanding of the VNS in wild rodents by providing a comprehensive characterization of the AOB using both routine histological techniques, such as Nissl staining, and more specific tools like immunohistochemical markers and lectin-histochemical staining.

### Histological study

One of the key findings of this study is the well-organized laminar structure of the AOB, which resembles that of the MOB but displays distinctive features. The arrangement of mitral cells in the AOB is less regular than in the MOB, as seen in other rodent species, such as hamsters (Nakajima et al. 1996), capybara (Torres et al. 2020) and octodon (Suárez and Mpodozis 2009). This irregular organization, combined with the fusion of the mitral and plexiform layers, likely reflects specific processing requirements for vomeronasal input, highlighting the functional specialization of the AOB in detecting chemosignals (Fernández-Aburto et al. 2020). The AOB of the fossorial water vole rat is also remarkable for the thickness of all its constitutive layers: nervous, glomerular, mitral-plexiform, and granular, as well as the lateral olfactory tract that traverses it. The mitral-plexiform layer is distinguished by the conspicuous number of principal cells that span the layer, consistently observed across the histological series from medial to lateral sections. Equally remarkable is the development of the mitral layer in the MOB, characterized by a high density of principal neurons with large somas. Previous research indicates that olfactory stimulation can enhance both the area and density of mitral cells (Johnson et al. 2013). Therefore, these morphological features are likely indicative of the heightened olfactory activity typical of this species.

An incidental finding in some of the studied individuals was the presence of parasitic cysts in the frontal lobe of the brain, and in some cases even in the MPL layer of the AOB. This is significant not only because it highlights the role of the species as a vector for diseases, such as toxoplasmosis, but also because these cysts can cause behavioral changes in the individual, leading to increased vulnerability to predators that are part of the parasites trophic chain (Vyas et al. 2007).

### Lectin histochemical study

The study of the olfactory bulb (OB) with lectins is of particular interest, as these markers enable selective characterization of the olfactory pathways—either the vomeronasal or the main olfactory pathway separately, or in conjunction— depending on the lectin used and the species involved (Park et al. 2014). Moreover, lectin histochemical labelling has facilitated the characterization of the anterior-posterior zonation of the AOB, which corresponds to the differential expression of G proteins ai2 and ao, which are respectively associated with the V1R and V2R vomeronasal chemoreceptor families (Jia and Halpern 1996). This zonation has been well characterized in the laboratory mouse (Salazar and Sánchez Quinteiro 2003) and rat (Sugai et al. 1997), and among wild rodents, in the hamster (Taniguchi et al. 1993b), capybara (Torres et al. 2020), degu (Suárez et al. 2011b), and beaver (Tomiyasu et al. 2022). In the case of the fossorial water vole, the validity of lectin labeling to reveal a clear antero-posterior zonation is confirmed. Out of the five lectins used, three of them, UEA, LEA, and STA, allowed its identification, whereas DBA and VVA, although they had different labeling patterns; DBA in superficial layers and VVA in the mitral-plexiform layer, in no case did they reveal the existence of zonation. These results coincide with the zonation observed with UEA in mice (Salazar et al. 2001; Barrios et al. 2014a; Kondoh et al. 2017), rats (Salazar and Sánchez Quinteiro 1998), and wallaby (Torres et al. 2022), with LEA in mice (Salazar et al. 2001), and STA in the hamster, although in the latter case it was much less noticeable (Taniguchi et al. 1993b).

Beyond zonation, each employed lectin presented unique patterns that were repeated in the AOB of all studied individuals. The only exception was UEA, a lectin specific to the L-fucose pathway, which, as characterized by Kondoh et al. (2017) in mice, shows a very variable expression pattern among individuals, independent of age and sex. Thus, depending on the individual studied, UEA in some instances was specific to the AOB, while in others it stained both the accessory and main olfactory bulbs to varying degrees. We have confirmed this observation in the study of the fossorial water vole, observing a similar variable pattern. Some authors suggest that these individual variations reflect rapid dynamic changes in the main and accessory olfactory bulbs, thereby causing alterations in marking patterns among individuals regardless of sex or age (Lipscomb et al. 2002; Kondoh et al. 2017). Furthermore, this individual variation may explain the disparate results found with UEA lectin in other rodents such as the rat where Barber (1989) and Ichikawa (1992), respectively considered UEA specific to the AOB or to the OB as a whole. A completely disparate case that exemplifies the great diversity in the expression of this lectin is the fact that UEA in rabbits does not recognize either the AOB or the MOB at all (Villamayor et al. 2020).

The study with LEA lectin exemplifies how special caution must be taken when assessing these individual differences. As a result of our histochemical staining along the entire series of the AOB in the sagittal plane, we found that the labeling pattern changes significantly. Depending on the area considered, areas are observed that are totally positive in the anterior-posterior axis. Others, mainly located in the central and lateral area of the BOA, show clear zonation. This process is a reflection of the asymmetries found in the compartmentalization of the accessory olfactory bulb in other rodents, such as the case of the degus (Suárez and Mpodozis 2009). Once again, this fact could explain the assumed absence of zonation with LEA lectin in the AOB of the rat (Salazar and Sánchez Quinteiro 1998) or rabbit (Villamayor et al. 2020) or the minimal differentiation observed by Taniguchi et al. in the hamster (1993b). STA lectin has been less studied in the BOA although the zonation observed by us in the water vole lends greater validity to the zonation detected by Ichikawa et al. in the rat (1992) and Taniguchi et al. in the hamster (1993b). DBA does not produce zonation but generates intense labelling in *Arvicola scherman*, which differentiates this species from the weak marking observed in mice (Salazar et al. 2001), in rats (Salazar and Sánchez Quinteiro 1998) and hamster (Taniguchi et al. 1993b). The peculiar labelling pattern produced by the VVA lectin in the fossorial water vole is striking, with strong positivity focused in the mitral plexiform layer that contrasts with the weak labelling observed in the mouse by Salazar et al. (2001) or in the rat by Takami et al. (1992). However, in this latter species, Salazar et al. (1998) and Ichikawa et al. (1992) found intense labelling in the AOB as did Shapiro et al. in the opossum (1995). Finally, in the case of the hamster, Taniguchi et al. (1993b) found a broad labelling pattern.

Our results confirm the utility of lectin markers in revealing distinct and variable patterns within both the AOB and the MOB, suggesting that lectin binding patterns can serve as differential markers to decode the intricate structural and functional organization within these regions. The differences in lectin binding among species suggest that these markers can be useful tools for a better understanding of the olfactory mechanisms across different groups of mammals.

### Immunohistochemical study

The immunohistochemical study with antibodies against the αo and αi2 subunits of G proteins is of particular interest, as the presence of both subunits is directly linked to the effective expression of the two main families of vomeronasal receptors, V1R and V2R. Specifically, Gαi2 is present in the transduction pathway of V1R receptors, while Gαo is involved in that of V2R. The immunohistochemical study in the AOB confirmed the segregated projection of both vomeronasal receptor families, with V1R receptors concentrated in the anterior zone of the AOB and V2R in the posterior zone. This pattern has also been observed in laboratory rodents such as rats (Shinohara et al. 1992) and mice (Jia and Halpern 1996), as well as in wild rodent species such as the beaver (Tomiyasu et al. 2022) and capybara (Torres et al. 2020). However, in the case of the squirrel, no similar zoning has been identified, as the only study conducted has found an AOB that exclusively expresses Gαi2 (Suárez et al. 2011a).

The anteroposterior zonation of the AOB has been correlated in laboratory rats (Matsuoka et al. 2001) and mice (Perez-Gomez et al. 2014) with the basal-apical zoning of the vomeronasal sensory epithelium. That is, neurons projecting to the anterior part of the AOB locate their somas in the apical part of the vomeronasal sensory neuroepithelium, while those projecting to the posterior zone are positioned basally. However, this basal-apical zonation has not been described in other rodents in which G protein expression in the VNO has been studied, such as in the case of the fossorial water vole (Ruiz-Rubio et al. 2024), the beaver (Tomiyasu et al. 2022), and the capybara (Torres et al. 2020), or even in mammals belonging to the segregated model, that is, those that exhibit anteroposterior zoning in the AOB, such as the rabbit (Torres et al. 2020) or the wallaby (Schneider et al. 2012). It is remarkably that the zonation found with lectins in the AOB of the fossorial water vole corresponds to the labelling pattern observed with UEA, LEA, and STA lectins. That is, the V1R axons reaching the anterior part of the AOB share a selective and analogous expression of glycoconjugates, which is different from that present in the V2R axons reaching the posterior part of the AOB.

Calcium-binding proteins exhibited in the AOB characteristic patterns indicating high activity in significant neuronal subpopulations within this region. Calbindin and calretinin revealed the laminated structure of the AOB. Specifically, both CB and CR produced dense and intense labeling in the nerve and glomerular layers of the AOB. In the mitral and granular layers, there was also intense labelling of the neuropil and cell bodies, although more pronounced in the case of calretinin. In the MOB, both antibodies produced a similar pattern, with dense labeling in the nerve and glomerular layers, but without staining the glomerular neuropil, concentrating exclusively in periglomerular cells. This pattern was similar to that described with anti-CR in the AOB and MOB of the rat (Jacobowitz and Winsky 1991). Differences in CB labeling were more pronounced across species within the superficial layers of the AOB compared to CR labeling. In particular, studies on rats (Porteros et al. 1995) and opossums (Jia and Halpern 2003; Jia and Halpern 2004) demonstrated that the neuropil displayed notably less staining than observed in fossorial water voles. Nonetheless, specific individual glomeruli in both rats and opossums displayed immunostaining and were associated with occasionally positively stained PG cells.

A remarkable finding was the identification with both anti-CB and anti-CR of a significant cluster of neurons located in the caudal subbulbar area. The role of these neuronal groups remains unknown, but their proximity to the AOB suggests they may be involved in the detection of chemical signals. To our knowledge, similar nuclei only have been described in the rat (Larriva-Sahd 2012), hamster (Ramón y Cajal 1902), rabbit (Young 1936; Villamayor et al. 2020), and hedgehog (Valverde et al. 1989), but in any case without the use of neuromarkers such as calbindin (CB) or calretinin (CR). Additionally, anti-calretinin strongly labelled the rostral migratory stream. Another calcium-binding protein studied was secretagogin, although in this case, the labeling pattern was more diffuse and less distinct, likely due to generalized labeling of the neuropil. Nevertheless, it was noticeable the presence of immunopositive periglomerular and mitral cells of the AOB.

The antibodies against MAP-2 and GAP-43 produced intense and revealing immunopositivity in the AOB of the fossorial water vole. Anti-MAP-2 is an excellent marker for the dendritic trees of mitral and granular cells (Dehmelt and Halpain 2005), but it does not stain their somas or axons (Bernhardt and Matus 1984). This is reflected in strong anti-MAP-2 immunolabelling in both the mitral-plexiform and granular layers, accompanied by immunonegativity in the somas of the mitral and granular cells of the AOB and LOT. Although MAP-2 immunolabeling should reveal the contribution of mitral cells to the glomeruli, as observed in the mouse AOB (Salazar et al. 2006), in the case of the fossorial water vole, this contribution is fainter.

GAP-43 serves as an effective marker for distinguishing between mature axons and regenerating nerve fibers, as its expression quickly diminishes in newly developed fibers once they have arrived at their destinations (Verhaagen et al. 1989; Ramakers et al. 1992). As it has been previously described in rabbits (Villamayor et al. 2020), the immunohistochemical labelling of GAP-43 in the fossorial water vole is a useful probe for discriminating the external and internal plexiform layers of the MOB, as well as the contribution of both plexiform layers to the mitral-plexiform layer of the AOB. According to GAP-43 immunostaining, we can conclude that the existence of two plexiform layers in the AOB is far from a theoretical concept.

The antibody against GAP-43 did not reveal the zonal organization of the AOB as it has been exceptionally demonstrated in rabbits (Villamayor et al. 2020). However, a remarkable observation -and to our knowledge, specific to the fossorial water vole-is the strong immunostaining of the somas of the mitral cells in the AOB, as well as the mitral and tufted cells of the MOB with anti-GAP-43. In adult neurons, high levels of GAP-43 expression are often linked to ongoing synaptic remodeling or heightened plasticity (Benowitz and Routtenberg 1997). In the case of the fossorial water vole -a subterranean animal that relies heavily on its sense of smell due to limited vision-the olfactory system may require enhanced adaptability to effectively process environmental cues (Fanjul et al. 2003). The increased GAP-43 expression in mitral and tufted cells could reflect a higher degree of synaptic plasticity necessary for refining olfactory and pheromonal processing under the unique conditions of an underground habitat. This adaptation might be crucial for activities such as foraging, navigation, and social interactions, where olfactory and semiochemical cues play a significant role. Further research would be necessary to confirm this hypothesis and to understand the specific roles of GAP-43 in the olfactory system of fossorial water voles.

Olfactory Marker Protein (OMP) is recognized as a specific marker for the superficial layers throughout the entire olfactory bulb. In fossorial water vole, OMP immunostaining is confined to the nervous strata and glomerular layers, exhibiting a slightly weaker intensity in the AOB compared to the main olfactory bulb MOB. The expression OMP is typical of mature neurons that carry olfactory and vomeronasal receptors (Rodewald et al. 2016; Nakamura et al. 2024). Although OMP expression has been widely studied across various mammalian VNOs (Smith et al. 2011; Dennis et al. 2020), investigations into its immunoreactivity within the AOB have been limited to certain species. These include rodents like mice, rats, and hamsters (Kream et al. 1984; Jia and Halpern 1996; Barrios et al. 2014a).

While the exact function of OMP remains to be fully elucidated, it is hypothesized to play a role in the maturation of olfactory and vomeronasal neurons (Bock et al. 2009). Moreover, OMP is implicated in the formation and refinement of the glomerular map, contributing to the precise organization of neural connections within the olfactory bulb (Albeanu et al. 2018). Correspondingly, OMP labeling has been observed to increase with age in both the VNO and the AOB (Verhaagen et al. 1989; Margolis et al. 1991).

Protein gene product 9.5 (PGP 9.5) is a soluble protein initially isolated from human brain tissue. It corresponds to the enzyme ubiquitin carboxyl-terminal hydrolase, which recycles ubiquitin from ubiquitin-linked protein complexes or polyubiquitin chains involved in proteolytic pathways (Hershko and Ciechanover 1992). Previous research has identified the expression of PGP 9.5 in both developing and mature olfactory and vomeronasal receptor cells of rats (Taniguchi et al. 1993a; Johnson et al. 1994). Moreover, studies have shown that PGP 9.5 predominantly localizes in the mitral and tufted cells within the main and accessory olfactory bulbs of rats and hamsters (Taniguchi et al. 1993a; Nakajima et al. 1996). In this study, we have confirmed the widespread immunoreactivity of PGP 9.5 in the accessory olfactory bulb of the fossorial water vole, with significant labelling in the mitral cells of the mitral-plexiform layer, as well as in periglomerular and short-axon cells within the glomerular layer. This distribution suggests that PGP 9.5 is essential for sustaining high levels of neuronal activity critical for olfactory processing

The immunohistochemical labeling with anti-SMI32 emphasizes the differences between the AOB and MOB in terms of neuronal activity and synaptic organization. The absence of anti-SMI32 labeling in the AOB, contrasted with strong labeling in the MOB, provides further evidence of the differential roles that these two structures play in olfactory signal processing. SMI32 is a marker for non-phosphorylated neurofilaments associated with mature neurons (Ang et al. 1991), which may indicate that the MOB handles more direct and rapid signal transmission, whereas the AOB might be involved in processing more complex, integrative signals related to social behaviors. Finally, the identification of active neurogenesis within the rostral migratory stream (RMS), immunolabelled with anti-DCX, is consistent with the known capacity for continuous neuronal turnover in the olfactory bulb (Huang et al. 2022). This reflects a dynamic process of neural regeneration in the olfactory system of the fossorial water vole, likely a crucial adaptation for maintaining olfactory sensitivity in the context of a changing subterranean environment.

In conclusion, our study underscores the structural and functional complexities of the AOB in rodents. The fossorial water vole, with its distinct reliance on chemical senses, provides an excellent model for understanding the neuroanatomical adaptations of the olfactory system in subterranean species. Future research should focus on exploring the functional consequences of the observed structural differences, particularly in the context of kairomonal and pheromonal detection, which could offer insights into the development of pest control strategies targeting chemical communication pathways (Fortes-Marco et al. 2015; Poissenot et al. 2023a; Poissenot et al. 2023b).

## AUTHORS CONTRIBUTION

Sara RUIZ-RUBIO: Conceptualization; Investigation; Funding acquisition; Writing - original draft. Irene ORTIZ-LEAL: Conceptualization; Investigation; Writing - review & editing; Methodology; Supervision. Mateo V. TORRES: Conceptualization; Investigation; Methodology; Supervision; Writing - review & editing. Mostafa G. A. ELSAYED: Conceptualization; Investigation; Methodology; Supervision; Writing - review & editing. Aitor SOMOANO: Investigation; Methodology; Supervision; Writing - review & editing. Pablo SANCHEZ-QUINTEIRO: Conceptualization; Investigation; Funding acquisition; Writing - original draft; Methodology; Writing - review & editing; Supervision.

## COMPLIANCE OF ETHICAL STANDARDS

### Ethical approval

Fossorial water voles are considered a key pest species in grasslands and their demographic densities need to be controlled in accordance with article 15 of the Law 43/2002 of plant health (BOE 2008) and therefore the practices undertaken in this study are considered as a recognised zootechnical purpose (Real Decreto 53/2013). Accordingly, ethics approval was not required for this study. The recommendations of the Directive of the European Parliament and the Council on the Protection of Animals Used for Scientific Purposes (Directive 2010/63/UE 2010) were considered in all procedures.

### Informed consent

No human subject was used in this study.

## ACKNOWLEDGEMENTS

The authors wish to thank the “Dirección Xeral de Gandaría, Agricultura e Industrias Agroalimentarias of the Consellería do Medio Rural of the XUNTA de GALICIA” for the financial, logistical support, and the trust placed in this project.

## REFERENCES

Airoldi, J.P. 1976. Le terrier de la forme fouisseuse du campagnol terrestre, Arvicola terrestris scherman Shaw (Mammalia, Rodentia). Z. Säugetierkd 41, pp. 23–42.

Albeanu, D.F., Provost, A.C., Agarwal, P., Soucy, E.R., Zak, J.D. and Murthy, V.N. 2018. Olfactory marker protein (OMP) regulates formation and refinement of the olfactory glomerular map. Nature Communications 9(1), p. 5073. doi: 10.1038/s41467-018-07544-9.

Ang, L.C., Munoz, D.G., Shul, D. and George, D.H. 1991. SMI-32 immunoreactivity in human striate cortex during postnatal development. Developmental Brain Research 61(1), pp. 103–109. doi: 10.1016/0165-3806(91)90119-4.

Apfelbach, R., Blanchard, C.D., Blanchard, R.J., Hayes, R.A. and McGregor, I.S. 2005. The effects of predator odors in mammalian prey species: A review of field and laboratory studies. Neuroscience & Biobehavioral Reviews 29(8), pp. 1123–1144. doi: 10.1016/j.neubiorev.2005.05.005.

Balmori-de La Puente, A., Ventura, J., Miñarro, M., Somoano, A., Hey, J. and Castresana, J. 2022. Divergence time estimation using ddRAD data and an isolation-with-migration model applied to water vole populations of Arvicola. Scientific Reports 12(1), p. 4065. doi: 10.1038/s41598-022-07877-y.

Barber, P.C. 1989. Ulex europeus agglutinin I binds exclusively to primary olfactory neurons in the rat nervous system. Neuroscience 30(1), pp. 1–9. doi: 10.1016/0306-4522(89)90348-5.

Barrios, A.W., Núñez, G., Sanchez Quinteiro, P. and Salazar, I. 2014a. Anatomy, histochemistry, and immunohistochemistry of the olfactory subsystems in mice. Frontiers in Neuroanatomy 8. Available at: http://journal.frontiersin.org/article/10.3389/fnana.2014.00063/abstract [Accessed: 11 December 2021].

Barrios, A.W., Sanchez Quinteiro, P. and Salazar, I. 2014b. The nasal cavity of the sheep and its olfactory sensory epithelium. Microsc. Res. Tech. 77(12), pp. 1052–1059. doi: 10.1002/jemt.22436.

Bembibre, C. and Strlič, M. 2022. From Smelly Buildings to the Scented Past: An Overview of Olfactory Heritage. Frontiers in Psychology 12, p. 718287. doi: 10.3389/fpsyg.2021.718287.

Benowitz, L.I. and Routtenberg, A. 1997. GAP-43: an intrinsic determinant of neuronal development and plasticity. Trends in Neurosciences 20(2), pp. 84–91. doi: 10.1016/S0166-2236(96)10072-2.

Bernhardt, R. and Matus, A. 1984. Light and electron microscopic studies of the distribution of microtubule-associated protein 2 in rat brain: A difference between dendritic and axonal cytoskeletons. The Journal of Comparative Neurology 226(2), pp. 203–221. doi: 10.1002/cne.902260205.

Bock, P., Rohn, K., Beineke, A., Baumgärtner, W. and Wewetzer, K. 2009. Site-specific population dynamics and variable olfactory marker protein expression in the postnatal canine olfactory epithelium. Journal of Anatomy 215(5), pp. 522–535. doi: 10.1111/j.1469-7580.2009.01147.x.

Boillat, M., Carleton, A. and Rodriguez, I. 2021. From immune to olfactory expression: neofunctionalization of formyl peptide receptors. Cell and Tissue Research 383(1), pp. 387–393. doi: 10.1007/s00441-020-03393-5.

Brennan, P.A. 2001. The vomeronasal system. Cellular and Molecular Life Sciences 58(4), pp. 546–555. doi: 10.1007/PL00000880.

Chengetanai, S., Bhagwandin, A., Bertelsen, M.F., Hård, T., Hof, P.R., Spocter, M.A. and Manger, P.R. 2020. The brain of the African wild dog. II. The olfactory system. Journal of Comparative Neurology 528(18), pp. 3285–3304. doi: 10.1002/cne.25007.

Chuah, M.I. and Zheng, D.R. 1987. Olfactory marker protein is present in olfactory receptor cells of human fetuses. Neuroscience 23(1), pp. 363–370. doi: 10.1016/0306-4522(87)90296-x.

Chun, J. et al. 2023. Glycoconjugate-Specific Developmental Changes in the Horse Vomeronasal Organ. Cells Tissues Organs, pp. 1–14. doi: 10.1159/000528883.

Dehmelt, L. and Halpain, S. 2005. The MAP2/Tau family of microtubule-associated proteins. Genome Biology 6(1), p. 204. doi: 10.1186/gb-2004-6-1-204.

Dennis, J.C., Stilwell, N.K., Smith, T.D., Park, T.J., Bhatnagar, K.P. and Morrison, E.E. 2020. Is the Mole Rat Vomeronasal Organ Functional? The Anatomical Record 303(2), pp. 318–329. doi: 10.1002/ar.24060.

Dielenberg, R.A. and McGregor, I.S. 2001. Defensive behavior in rats towards predatory odors: a review. Neuroscience & Biobehavioral Reviews 25(7–8), pp. 597–609. doi: 10.1016/S0149-7634(01)00044-6.

Dulac, C. and Axel, R. 1995. A novel family of genes encoding putative pheromone receptors in mammals. Cell 83(2), pp. 195–206. doi: 10.1016/0092-8674(95)90161-2.

Ennis, M. and Holy, T.E. 2015. Anatomy and Neurobiology of the Main and Accessory Olfactory Bulbs. In: Doty, R. L. ed. Handbook of Olfaction and Gustation. Hoboken, NJ, USA: John Wiley & Sons, Inc, pp. 157–182. Available at: https://onlinelibrary.wiley.com/doi/10.1002/9781118971758.ch8 [Accessed: 12 December 2021].

Espí, A., Del Cerro, A., Somoano, A., García, V., M. Prieto, J., Barandika, J.F. and García-Pérez, A.L. 2017. Borrelia burgdorferi sensu lato prevalence and diversity in ticks and small mammals in a Lyme borreliosis endemic Nature Reserve in North-Western Spain. Incidence in surrounding human populations. Enfermedades Infecciosas y Microbiología Clínica 35(9), pp. 563–568. doi: 10.1016/j.eimc.2016.06.011.

Fanjul, M.S., Zenuto, R.R. and Busch, C. 2003. Use of olfaction for sexual recognition in the subterranean rodentCtenomys talarum. Acta Theriologica 48(1), pp. 35–46. doi: 10.1007/BF03194264.

Fernández-Aburto, P., Delgado, S.E., Sobrero, R. and Mpodozis, J. 2020. Can social behaviour drive accessory olfactory bulb asymmetries? Sister species of caviomorph rodents as a case in point. Journal of Anatomy 236(4), pp. 612–621. doi: 10.1111/joa.13126.

Fortes-Marco, L., Lanuza, E. and Martinez-Garcia, F. 2013. Of Pheromones and Kairomones: What Receptors Mediate Innate Emotional Responses?: Pheromones and Kairomones. The Anatomical Record 296(9), pp. 1346–1363. doi: 10.1002/ar.22745.

Fortes-Marco, L., Lanuza, E., Martínez-García, F. and Agustín-Pavón, C. 2015. Avoidance and contextual learning induced by a kairomone, a pheromone and a common odorant in female CD1 mice. Frontiers in Neuroscience 9. Available at: http://journal.frontiersin.org/Article/10.3389/fnins.2015.00336/abstract [Accessed: 15 September 2022].

Frahm, H.D. and Bhatnagar, K.P. 1980. Comparative morphology of the accessory olfactory bulb in bats. Journal of Anatomy 130(Pt 2), pp. 349–365.

Franceschini, V., Lazzari, M., Revoltella, R.P. and Ciani, F. 1994. Histochemical study by lectin binding of surface glycoconjugates in the developing olfactory system of rat. International Journal of Developmental Neuroscience: The Official Journal of the International Society for Developmental Neuroscience 12(3), pp. 197–206. doi: 10.1016/0736-5748(94)90041-8.

Fuehrer, H.-P., Blöschl, I., Siehs, C. and Hassl, A. 2010. Detection of Toxoplasma gondii, Neospora caninum, and Encephalitozoon cuniculi in the brains of common voles (Microtus arvalis) and water voles (Arvicola terrestris) by gene amplification techniques in western Austria (Vorarlberg). Parasitology Research 107(2), pp. 469–473. doi: 10.1007/s00436-010-1905-z.

Giraudoux, P., Charbonell, N., Deter, J., Chaval, Y., Cosson, J.F. and Raoul, F. 2009. Maladies transmissibles à l’homme. In: Le campagnol terrestre. Prévention et contrôle des populations. Delattre P. & Giraudoux P. Versailles Cedex: Éditions Quæ, pp. 101–110.

Halpern, M. and Martínez-Marcos, A. 2003. Structure and function of the vomeronasal system: an update. Progress in Neurobiology 70(3), pp. 245–318. doi: 10.1016/S0301-0082(03)00103-5.

Hershko, A. and Ciechanover, A. 1992. The ubiquitin system for protein degradation. Annual Review of Biochemistry 61, pp. 761–807. doi: 10.1146/annurev.bi.61.070192.003553.

Holy, T.E. 2018. The Accessory Olfactory System: Innately Specialized or Microcosm of Mammalian Circuitry? Annual Review of Neuroscience 41(1), pp. 501–525. doi: 10.1146/annurev-neuro-080317-061916.

Horii, Y., Nikaido, Y., Nagai, K. and Nakashima, T. 2010. Exposure to TMT odor affects adrenal sympathetic nerve activity and behavioral consequences in rats. Behavioural Brain Research 214(2), pp. 317–322. doi: 10.1016/j.bbr.2010.05.047.

Huang, J.S. et al. 2022. Immature olfactory sensory neurons provide behaviourally relevant sensory input to the olfactory bulb. Nature Communications 13(1), p. 6194. doi: 10.1038/s41467-022-33967-6.

Ichikawa, M., Osada, T. and Ikai, A. 1992. Bandeiraea simplicifolia lectin I and Vicia villosa agglutinin bind specifically to the vomeronasal axons in the accessory olfactory bulb of the rat. Neuroscience Research 13(1), pp. 73–79. doi: 10.1016/0168-0102(92)90035-B.

Jacobowitz, D.M. and Winsky, L. 1991. Immunocytochemical localization of calretinin in the forebrain of the rat. The Journal of Comparative Neurology 304(2), pp. 198–218. doi: 10.1002/cne.903040205.

Jia, C. and Halpern, M. 1996. Subclasses of vomeronasal receptor neurons: differential expression of G proteins (Giα2 and Goα) and segregated projections to the accessory olfactory bulb. Brain Research 719(1–2), pp. 117–128. doi: 10.1016/0006-8993(96)00110-2.

Jia, C. and Halpern, M. 2003. Calbindin D28K immunoreactive neurons in vomeronasal organ and their projections to the accessory olfactory bulb in the rat. Brain Research 977(2), pp. 261–269. doi: 10.1016/S0006-8993(03)02693-3.

Jia, C. and Halpern, M. 2004. Calbindin D28k, parvalbumin, and calretinin immunoreactivity in the main and accessory olfactory bulbs of the gray short-tailed opossum,Monodelphis domestica. Journal of Morphology 259(3), pp. 271–280. doi: 10.1002/jmor.10166.

Johnson, E.W., Eller, P.M. and Jafek, B.W. 1994. Protein gene product 9.5 in the developing and mature rat vomeronasal organ. Developmental Brain Research 78(2), pp. 259–264. doi: 10.1016/0165-3806(94)90034-5.

Johnson, G. and Jope, R. 1992. The role of microtubule-associated protein 2 (MAP-2) in neuronal growth, plasticity, and degeneration. Journal of Neuroscience Research 33(4), pp. 505–512. doi: 10.1002/jnr.490330402.

Johnson, M.C., Biju, K.C., Hoffman, J. and Fadool, D.A. 2013. Odor Enrichment Sculpts the Abundance of Olfactory Bulb Mitral Cells. Neuroscience letters 0, pp. 173–178. doi: 10.1016/j.neulet.2013.02.027.

Keller, L.-A., Niedermeier, S., Claassen, L. and Popp, A. 2022. Comparative lectin histochemistry on the murine respiratory tract and primary olfactory pathway using a fully automated staining procedure. Acta Histochemica 124(3), p. 151877. doi: 10.1016/j.acthis.2022.151877.

Kelliher, K.R., Baum, M.J. and Meredith, M. 2001. The Ferret’s vomeronasal organ and accessory olfactory bulb: Effect of hormone manipulation in adult males and females. The Anatomical Record 263(3), pp. 280–288. doi: 10.1002/ar.1097.

Kondoh, D., Kamikawa, A., Sasaki, M. and Kitamura, N. 2017. Localization of α1-2 Fucose Glycan in the Mouse Olfactory Pathway. Cells Tissues Organs 203(1), pp. 20–28. doi: 10.1159/000447009.

Kondoh, D., Kawai, Y.K., Watanabe, K. and Muranishi, Y. 2022. Artiodactyl livestock species have a uniform vomeronasal system with a vomeronasal type 1 receptor (V1R) pathway. Tissue and Cell 77, p. 101863. doi: 10.1016/j.tice.2022.101863.

Kream, R.M., Davis, B.J., Kawano, T., Margolis, F.L. and Macrides, F. 1984. Substance P and catecholaminergic expression in neurons of the hamster main olfactory bulb. The Journal of Comparative Neurology 222(1), pp. 140–154. doi: 10.1002/cne.902220112.

Larriva-Sahd, J. 2008. The accessory olfactory bulb in the adult rat: A cytological study of its cell types, neuropil, neuronal modules, and interactions with the main olfactory system. The Journal of Comparative Neurology 510(3), pp. 309–350. doi: 10.1002/cne.21790.

Larriva-Sahd, J. 2012. Cytological organization of the alpha component of the anterior olfactory nucleus and olfactory limbus. Frontiers in Neuroanatomy 6. Available at: http://journal.frontiersin.org/article/10.3389/fnana.2012.00023/abstract [Accessed: 27 December 2021].

Lee, V.M., Otvos, L., Carden, M.J., Hollosi, M., Dietzschold, B. and Lazzarini, R.A. 1988. Identification of the major multiphosphorylation site in mammalian neurofilaments. Proceedings of the National Academy of Sciences 85(6), pp. 1998–2002. doi: 10.1073/pnas.85.6.1998.

Lipscomb, B., Treloar, H. and Greer, C. 2002. Cell surface carbohydrates reveal heterogeneity in olfactory receptor cell axons in the mouse. Cell and Tissue Research 308(1), pp. 7–17. doi: 10.1007/s00441-002-0532-0.

Lis, H. and Sharon, N. 1998. Lectins: Carbohydrate-Specific Proteins That Mediate Cellular Recognition. Chemical Reviews 98(2), pp. 637–674. doi: 10.1021/cr940413g.

Margolis, F.L., Verhaagen, J., Biffo, S., Huang, F.L. and Grillo, M. 1991. Regulation of gene expression in the olfactory neuroepithelium: a neurogenetic matrix. Progress in Brain Research 89, pp. 97–122. doi: 10.1016/s0079-6123(08)61718-5.

Martín-López, E., Corona, R. and López-Mascaraque, L. 2012. Postnatal characterization of cells in the accessory olfactory bulb of wild type and reeler mice. Frontiers in Neuroanatomy 6. Available at: http://journal.frontiersin.org/article/10.3389/fnana.2012.00015/abstract [Accessed: 22 August 2023].

Matsuoka, M., Yoshida-Matsuoka, J., Iwasaki, N., Norita, M., Costanzo, R.M. and Ichikawa, M. 2001. Immunocytochemical Study of Gi2α and Goα on the Epithelium Surface of the Rat Vomeronasal Organ. Chemical Senses 26(2), pp. 161–166. doi: 10.1093/chemse/26.2.161.

Meisami, E. and Bhatnagar, K.P. 1998. Structure and diversity in mammalian accessory olfactory bulb. Microscopy Research and Technique 43(6), pp. 476–499. doi: 10.1002/(SICI)1097-0029(19981215)43:6<476::AID-JEMT2>3.0.CO;2-V.

Menini, A. ed. 2010. The Neurobiology of Olfaction. Boca Raton (FL): CRC Press/Taylor & Francis. Available at: http://www.ncbi.nlm.nih.gov/books/NBK55980/ [Accessed: 9 September 2024].

Mucignat, C. 2004. High-resolution Magnetic Resonance Spectroscopy of the Mouse Vomeronasal Organ. Chemical Senses 29(8), pp. 693–696. doi: 10.1093/chemse/bjh073.

Nagnan-Le Meillour, P. et al. 2019. Identification of potential chemosignals in the European water vole Arvicola terrestris. Scientific Reports 9(1), p. 18378. doi: 10.1038/s41598-019-54935-z.

Nakajima, T., Okamura, M., Ogawa, K. and Taniguchi, K. 1996. Immunohistochemical and enzyme histochemical characteristics of short axon cells in the olfactory bulb of the golden hamster. The Journal of Veterinary Medical Science 58(9), pp. 903–908. doi: 10.1292/jvms.58.903.

Nakajima, T., Sakaue, M., Kato, M., Saito, S., Ogawa, K. and Taniguchi, K. 1998. Immunohistochemical and enzyme-histochemical study on the accessory olfactory bulb of the dog. The Anatomical Record 252(3), pp. 393–402. doi: 10.1002/(SICI)1097-0185(199811)252:3<393::AID-AR7>3.0.CO;2-T.

Nakamura, Y., Miwa, T., Shiga, H., Sakata, H., Shigeta, D. and Hatta, T. 2024. Histological changes in the olfactory bulb and rostral migratory stream due to interruption of olfactory input. Auris, Nasus, Larynx 51(3), pp. 517–524. doi: 10.1016/j.anl.2024.01.009.

Ortiz-Leal, I., Torres, M.V., Barreiro-Vázquez, J., López-Beceiro, A., Fidalgo, L., Shin, T. and Sanchez-Quinteiro, P. 2024. The vomeronasal system of the wolf (*Canis lupus signatus*): The singularities of a wild canid. Journal of Anatomy 00(1–28), p. joa.14024. doi: 10.1111/joa.14024.

Ortiz-Leal, I., Torres, M.V., López-Beceiro, A., Fidalgo, L., Shin, T. and Sanchez-Quinteiro, P. 2024. First Immunohistochemical Demonstration of the Expression of a Type-2 Vomeronasal Receptor, V2R2, in Wild Canids. International Journal of Molecular Sciences 25(13), p. 7291. doi: 10.3390/ijms25137291.

Ortiz-Leal, I., Torres, M.V., López-Callejo, L.N., Fidalgo, L.E., López-Beceiro, A. and Sanchez-Quinteiro, P. 2022a. Comparative Neuroanatomical Study of the Main Olfactory Bulb in Domestic and Wild Canids: Dog, Wolf and Red Fox. Animals 12(9), p. 1079. doi: 10.3390/ani12091079.

Ortiz-Leal, I., Torres, M.V., Vargas-Barroso, V., Fidalgo, L.E., López-Beceiro, A.M., Larriva-Sahd, J.A. and Sánchez-Quinteiro, P. 2023. The olfactory limbus of the red fox (Vulpes vulpes). New insights regarding a noncanonical olfactory bulb pathway. Frontiers in Neuroanatomy 16, p. 1097467. doi: 10.3389/fnana.2022.1097467.

Ortiz-Leal, I., Torres, M.V., Villamayor, P.R., Fidalgo, L.E., López-Beceiro, A. and Sanchez-Quinteiro, P. 2022b. Can domestication shape Canidae brain morphology? The accessory olfactory bulb of the red fox as a case in point. Annals of Anatomy - Anatomischer Anzeiger 240, p. 151881. doi: 10.1016/j.aanat.2021.151881.

Ortiz-Leal, I., Torres, M.V., Villamayor, P.R., López-Beceiro, A. and Sanchez-Quinteiro, P. 2020. The vomeronasal organ of wild canids: the fox (*Vulpes vulpes*) as a model. Journal of Anatomy 237(5), pp. 890–906. doi: 10.1111/joa.13254.

Overath, P., Sturm, T. and Rammensee, H.-G. 2014. Of volatiles and peptides: in search for MHC-dependent olfactory signals in social communication. Cellular and Molecular Life Sciences 71(13), pp. 2429–2442. doi: 10.1007/s00018-014-1559-6.

Papes, F., Logan, D.W. and Stowers, L. 2010. The Vomeronasal Organ Mediates Interspecies Defensive Behaviors through Detection of Protein Pheromone Homologs. Cell 141(4), pp. 692–703. doi: 10.1016/j.cell.2010.03.037.

Park, C. et al. 2014. A morphological study of the vomeronasal organ and the accessory olfactory bulb in the Korean roe deer, Capreolus pygargus. Acta Histochemica 116(1), pp. 258–264. doi: 10.1016/j.acthis.2013.08.003.

Perez-Gomez, A., Stein, B., Leinders-Zufall, T. and Chamero, P. 2014. Signaling mechanisms and behavioral function of the mouse Basal vomeronasal neuroepithelium. Frontiers in neuroanatomy 8, p. 135. doi: 10.3389/fnana.2014.00135.

Poissenot, K. et al. 2023a. Sexual discrimination and attraction through scents in the water vole, Arvicola terrestris. Journal of Comparative Physiology A. Available at: https://link.springer.com/10.1007/s00359-023-01671-5 [Accessed: 11 September 2023].

Poissenot, K., Porte, C., Chesneau, D. and Keller, M. 2023b. Exploration of Olfactory Communication in the Water Vole, Arvicola terrestris. In: Schaal, B., Rekow, D., Keller, M., and Damon, F. eds. Chemical Signals in Vertebrates 15. Cham: Springer International Publishing, pp. 153–163. Available at: https://link.springer.com/10.1007/978-3-031-35159-4_8 [Accessed: 26 November 2023].

Porteros, A., Arévalo, R., Crespo, C., García-Ojeda, E., Briñón, J.G., Aijón, J. and Alonso, J.R. 1995. Calbindin D-28k immunoreactivity in the rat accessory olfactory bulb. Brain Research 689(1), pp. 93–100. doi: 10.1016/0006-8993(95)00547-4.

Ramakers, G.J.A., Verhaagen, J., Oestreicher, A.B., Margolis, F.L., Van Bergen En Henegouwen, P.M.P. and Gispen, W.H. 1992. Immunolocalization of B-50 (GAP-43) in the mouse olfactory bulb: Predominant presence in preterminal axons. Journal of Neurocytology 21(12), pp. 853–869. doi: 10.1007/BF01191683.

Ramón y Cajal, S.R. 1902. Textura del lobulo olfativo accesorio. 1, pp. 141–150.

Robardet, E., Giraudoux, P., Caillot, C., Augot, D., Boue, F. and Barrat, J. 2011. Fox defecation behaviour in relation to spatial distribution of voles in an urbanised area: An increasing risk of transmission of Echinococcus multilocularis? International Journal for Parasitology 41(2), pp. 145–154. doi: 10.1016/j.ijpara.2010.08.007.

Rodewald, A., Gisder, D., Gebhart, V.M., Oehring, H. and Jirikowski, G.F. 2016. Distribution of olfactory marker protein in the rat vomeronasal organ. Journal of Chemical Neuroanatomy 77, pp. 19–23. doi: 10.1016/j.jchemneu.2016.04.002.

Ruiz-Rubio, S., Ortiz-Leal, I., Torres, M.V., Somoano, A. and Sanchez-Quinteiro, P. 2024. Do fossorial water voles have a functional vomeronasal organ? A histological and immunohistochemical study. Anatomical Record (Hoboken, N.J.: 2007) 307(8), pp. 2912–2932. doi: 10.1002/ar.25374.

Ryba, N.J.P. and Tirindelli, R. 1997. A New Multigene Family of Putative Pheromone Receptors. Neuron 19(2), pp. 371–379. doi: 10.1016/S0896-6273(00)80946-0.

Saito, S., Nii, Y. and Taniguchi, K. 1999. Heterogeneous expression of glycoconjugates among individual glomeruli of the hamster main olfactory bulb. Chemical Senses 24(5), pp. 509–515. doi: 10.1093/chemse/24.5.509.

Salazar, I., Barrios, A.W. and Sánchez-Quinteiro, P. 2016. Revisiting the Vomeronasal System From an Integrated Perspective. The Anatomical Record 299(11), pp. 1488–1491. doi: 10.1002/ar.23470.

Salazar, I., Lombardero, M., Cifuentes, J.M., Quinteiro, P.S. and Aleman, N. 2003. Morphogenesis and growth of the soft tissue and cartilage of the vomeronasal organ in pigs. Journal of Anatomy 202(6), pp. 503–514. doi: 10.1046/j.1469-7580.2003.00183.x.

Salazar, I. and Sánchez Quinteiro, P. 1998. Lectin binding patterns in the vomeronasal organ and accessory olfactory bulb of the rat. Anatomy and Embryology 198(4), pp. 331–339. doi: 10.1007/s004290050188.

Salazar, I. and Sánchez Quinteiro, P. 2003. Differential development of binding sites for four lectins in the vomeronasal system of juvenile mouse: from the sensory transduction site to the first relay stage. Brain Research 979(1–2), pp. 15–26. doi: 10.1016/S0006-8993(03)02835-X.

Salazar, I., Sánchez Quinteiro, P., Cifuentes, J.M., Fernández, P. and Lombardero, M. 1997. Distribution of the arterial supply to the vomeronasal organ in the cat. The Anatomical Record 247(1), pp. 129–136. doi: 10.1002/(SICI)1097-0185(199701)247:1<129::AID-AR15>3.0.CO;2-R.

Salazar, I., Sanchez-Quinteiro, P., Cifuentes, J.M. and De Troconiz, P.F. 2006. General organization of the perinatal and adult accessory olfactory bulb in mice. The Anatomical Record Part A: Discoveries in Molecular, Cellular, and Evolutionary Biology 288A(9), pp. 1009–1025. doi: 10.1002/ar.a.20366.

Salazar, I., Sanchez-Quinteiro, P., Lombardero, M. and Cifuentes, J.M. 2001. Histochemical Identification of Carbohydrate Moieties in the Accessory Olfactory Bulb of the Mouse Using a Panel of Lectins. Chemical Senses 26(6), pp. 645–652. doi: 10.1093/chemse/26.6.645.

Sbarbati, A. and Osculati, F. 2006. Allelochemical Communication in Vertebrates: Kairomones, Allomones and Synomones. Cells Tissues Organs 183(4), pp. 206–219. doi: 10.1159/000096511.

Scalia, F. and Winans, S.S. 1975. The differential projections of the olfactory bulb and accessory olfactory bulb in mammals. The Journal of Comparative Neurology 161(1), pp. 31–55. doi: 10.1002/cne.901610105.

Schneider, N.Y., Fletcher, T.P., Shaw, G. and Renfree, M.B. 2012. Goα Expression in the Vomeronasal Organ and Olfactory Bulb of the Tammar Wallaby. Chemical Senses 37(6), pp. 567–577. doi: 10.1093/chemse/bjs040.

Schröder, H., Moser, N. and Huggenberger, S. 2020. The Mouse Olfactory System. In: Schröder, H., Moser, N., and Huggenberger, S. eds. Neuroanatomy of the Mouse: An Introduction. Cham: Springer International Publishing, pp. 319–331. Available at: 10.1007/978-3-030-19898-5_14 [Accessed: 25 September 2022].

Scott, K. 2003. Sex and the MHC. Developmental Cell 4(3), pp. 290–291. doi: 10.1016/S1534-5807(03)00066-2.

Shapiro, L.S., Halpern, M. and Ee, P.-L. 1995. Lectin histochemical identification of carbohydrate moieties in opossum chemosensory systems during development, with special emphasis on VVA-identified subdivisions in the accessory olfactory bulb. Journal of Morphology 224(3), pp. 331–349. doi: 10.1002/jmor.1052240307.

Shin, T., Kim, J., Choi, Y. and Ahn, M. 2017. Glycan diversity in the vomeronasal organ of the Korean roe deer, Capreolus pygargus : A lectin histochemical study. Acta Histochemica 119(8), pp. 778–785. doi: 10.1016/j.acthis.2017.10.001.

Shinohara, H., Asano, T. and Kato, K. 1992. Differential localization of G-proteins Gi and Go in the accessory olfactory bulb of the rat. The Journal of Neuroscience 12(4), pp. 1275–1279. doi: 10.1523/JNEUROSCI.12-04-01275.1992.

Smith, T.D., Dennis, J.C., Bhatnagar, K.P., Garrett, E.C., Bonar, C.J. and Morrison, E.E. 2011. Olfactory marker protein expression in the vomeronasal neuroepithelium of tamarins (Saguinus spp). Brain Research 1375, pp. 7–18. doi: 10.1016/j.brainres.2010.12.069.

Smithson, L.J. and Kawaja, M.D. 2009. A comparative examination of biomarkers for olfactory ensheathing cells in cats and guinea pigs. Brain Research 1284, pp. 41–53. doi: 10.1016/j.brainres.2009.06.011.

Somoano, A. 2020. The role of the montane water vole (Arvicola scherman) as a crop pest in NW Spain: since when? *Galemys*, Spanish Journal of Mammalogy 32, pp. 61–63. doi: 10.7325/Galemys.2020.N1.

Suárez, R., Fernández-Aburto, P., Manger, P.R. and Mpodozis, J. 2011a. Deterioration of the Gαo Vomeronasal Pathway in Sexually Dimorphic Mammals. Callaerts, P. ed. PLoS ONE 6(10), p. e26436. doi: 10.1371/journal.pone.0026436.

Suárez, R. and Mpodozis, J. 2009. Heterogeneities of size and sexual dimorphism between the subdomains of the lateral-innervated accessory olfactory bulb (AOB) of *Octodon degus* (Rodentia: Hystricognathi). Behavioural Brain Research 198(2), pp. 306–312. doi: 10.1016/j.bbr.2008.11.009.

Suárez, R., Santibáñez, R., Parra, D., Coppi, A.A., Abrahão, L.M.B., Sasahara, T.H.C. and Mpodozis, J. 2011b. Shared and differential traits in the accessory olfactory bulb of caviomorph rodents with particular reference to the semiaquatic capybara: The AOB of capybaras and other caviomorphs. Journal of Anatomy 218(5), pp. 558–565. doi: 10.1111/j.1469-7580.2011.01357.x.

Sugai, T., Sugitani, M. and Onoda, N. 1997. Subdivisions of the guinea-pig accessory olfactory bulb revealed by the combined method with immunohistochemistry, electrophysiological, and optical recordings. Neuroscience 79(3), pp. 871–885. doi: 10.1016/S0306-4522(96)00690-2.

Switzer, III., R.C., Johnson, J.I. and Kirsch, J.A.W. 1980. Phylogeny Through Brain Traits. Brain, Behavior and Evolution 17(5), pp. 339–363. doi: 10.1159/000121808.

Takahashi, L.K. 2014. Olfactory systems and neural circuits that modulate predator odor fear. Frontiers in Behavioral Neuroscience 8. Available at: http://journal.frontiersin.org/article/10.3389/fnbeh.2014.00072/abstract [Accessed: 8 September 2023].

Takami, S., Graziadei, P.P.C. and Ichikawa, M. 1992. The differential staining patterns of two lectins in the accessory olfactory bulb of the rat. Brain Research 598(1– 2), pp. 337–342. doi: 10.1016/0006-8993(92)90204-M.

Taniguchi, K., Harumi, S., Okamura, M. and Ogawa, K. 1993a. Immunohistochemical demonstration of protein gene product 9.5 (PGP 9.5) in the primary olfactory system of the rat. Neuroscience Letters 156(1–2), pp. 24–26. doi: 10.1016/0304-3940(93)90430-S.

Taniguchi, K., Nii, Y. and Ogawa, K. 1993b. Subdivisions of the accessory olfactory bulb, as demonstrated by lectin-histochemistry in the golden hamster. Neuroscience Letters 158(2), pp. 185–188. doi: 10.1016/0304-3940(93)90260-R.

Taroc, E.Z.M. et al. 2020. Gli3 Regulates Vomeronasal Neurogenesis, Olfactory Ensheathing Cell Formation, and GnRH-1 Neuronal Migration. The Journal of Neuroscience 40(2), pp. 311–326. doi: 10.1523/JNEUROSCI.1977-19.2019.

Tomiyasu, J., Kondoh, D., Sakamoto, H., Matsumoto, N., Haneda, S. and Matsui, M. 2018. Lectin histochemical studies on the olfactory gland and two types of gland in vomeronasal organ of the brown bear. Acta Histochemica 120(6), pp. 566–571. doi: 10.1016/j.acthis.2018.07.003.

Tomiyasu, J., Korzekwa, A., Kawai, Y.K., Robstad, C.A., Rosell, F. and Kondoh, D. 2022. The vomeronasal system in semiaquatic beavers. Journal of Anatomy 241(3), pp. 809–819. doi: 10.1111/joa.13671.

Torres, M.V., Ortiz-Leal, I., Ferreiro, A., Rois, J.L. and Sanchez-Quinteiro, P. 2021. Neuroanatomical and Immunohistological Study of the Main and Accessory Olfactory Bulbs of the Meerkat (*Suricata suricatta*). Animals 12(1), p. 91. doi: 10.3390/ani12010091.

Torres, M.V., Ortiz-Leal, I., Ferreiro, A., Rois, J.L. and Sanchez-Quinteiro, P. 2023. Immunohistological study of the unexplored vomeronasal organ of an endangered mammal, the dama gazelle (*Nanger dama*). Microscopy Research and Technique 86(9), pp. 1206–1233. doi: 10.1002/jemt.24392.

Torres, M.V., Ortiz-Leal, I., Villamayor, P.R., Ferreiro, A., Rois, J.L. and Sanchez-Quinteiro, P. 2020. The vomeronasal system of the newborn capybara: a morphological and immunohistochemical study. Scientific Reports 10(1), p. 13304. doi: 10.1038/s41598-020-69994-w.

Torres, M.V., Ortiz-Leal, I., Villamayor, P.R., Ferreiro, A., Rois, J.L. and Sanchez-Quinteiro, P. 2022. Does a third intermediate model for the vomeronasal processing of information exist? Insights from the macropodid neuroanatomy. Brain Structure and Function 227(3), pp. 881–899. doi: 10.1007/s00429-021-02425-2.

Valverde, F., López-Mascaraque, L. and De Carlos, J.A. 1989. Structure of the nucleus olfactorius anterior of the hedgehog (*Erinaceus europaeus*). Journal of Comparative Neurology 279(4), pp. 581–600. doi: 10.1002/cne.902790407.

Verhaagen, J., Oestreicher, A., Gispen, W. and Margolis, F. 1989. The expression of the growth associated protein B50/GAP43 in the olfactory system of neonatal and adult rats. The Journal of Neuroscience 9(2), pp. 683–691. doi: 10.1523/JNEUROSCI.09-02-00683.1989.

Villamayor, P.R., Cifuentes, J.M., Quintela, L., Barcia, R. and Sanchez-Quinteiro, P. 2020. Structural, morphometric and immunohistochemical study of the rabbit accessory olfactory bulb. Brain Structure and Function 225(1), pp. 203–226. doi: 10.1007/s00429-019-01997-4.

Vyas, A., Kim, S.-K. and Sapolsky, R.M. 2007. The effects of Toxoplasma infection on rodent behavior are dependent on dose of the stimulus. Neuroscience 148(2), pp. 342–348. doi: 10.1016/j.neuroscience.2007.06.021.

Winans, S.S. and Scalia, F. 1970. Amygdaloid Nucleus: New Afferent Input from the Vomeronasal Organ. Science 170(3955), pp. 330–332. doi: 10.1126/science.170.3955.330.

Yohe, L.R. and Krell, N.T. 2023. An updated synthesis of and outstanding questions in the olfactory and vomeronasal systems in bats: Genetics asks questions only anatomy can answer. The Anatomical Record, p. ar.25290. doi: 10.1002/ar.25290.

Young, M.W. 1936. The nuclear pattern and fiber connections of the non-cortical centers of the telencephalon of the rabbit (Lepus cuniculus). Journal of Comparative Neurology 65(1), pp. 295–401. doi: 10.1002/cne.900650112.

